# Predicting Developmental Norms from Baseline Cortical Thickness in Longitudinal Studies

**DOI:** 10.1101/2025.07.11.664301

**Authors:** Philipp Seidel, Tobias Kaufmann, Thomas Wolfers

## Abstract

**Background:** Normative models have gained popularity in computational psychiatry for studying individual-level differences relative to population norms in biological data such as brain imaging, where measures like cortical thickness are typically predicted from variables such as age and sex. Nearly all published models to date are based on cross-sectional data, limiting their ability to predict longitudinal change.

**Methods:** Here, we used longitudinal brain data from the Adolescent Brain Cognitive Development (ABCD) study, comprising cortical thickness measures from 180 regions per hemisphere in youths at baseline (N=6179; 47% females), 2-year (N=6179; 47% females), and 4-year (N=805; 45% females) follow-up. A training set was established from baseline and 2-year follow-up data (N=5374; 47% females), while data from individuals with all three time points available served as an independent test set (N=805; 45% females). We developed sex-specific Baseline-Conditioned Norms (B-Norms) that predict brain region thickness at follow-up based on baseline thickness, baseline age, and follow-up age, and compared them to sex-specific Cross-Sectional Norms (C-Norms) based on age alone.

**Results:** Out-of-sample testing in 2-year and 4-year follow-up data showed that B-Norms consistently provided better fits than C-Norms for nearly all cortical regions. Explained variance was higher in B-Norms than in C-Norms. No significant differences between time points (p = 0.45) were detected. Repeated measures ANOVA revealed differences in higher-order moments (e.g., skewness and kurtosis) for both models; for example, skewness varied by model, sex, time point, and their interactions.

**Conclusion:** While improved fit alone does not necessarily indicate a superior normative model, since normative models aim to capture population variance rather than simply optimize fit, we demonstrated that four regions were associated with pubertal changes in B-Norms but not in C-Norms, suggesting enhanced sensitivity of B-Norms to developmental processes. Together, our findings highlight the potential of B-Norms for capturing normative variation in longitudinal structural brain change.

**Highlights:**

- Baseline-Conditioned (B-Norm) models consistently outperform Cross-Sectional (C-Norm) models across all cortical regions.
- B-Norm deviation scores show stronger associations with pubertal progression in females than C-Norm deviations, indicating higher developmental sensitivity.
- This contrasts with brain-age models, where improved fit typically reduces the association between residuals and downstream clinical or developmental variation.

**Summary:** Normative modeling has been applied to study how brain measures, such as gray matter thickness or volume, change across development. These models help identify how an individual’s brain may differ from what is typical for their age or sex, which could eventually support more personalized treatments.

However, most existing models use only one-time (cross-sectional) data, meaning they cannot capture how the brain changes over time. Longitudinal data, tracking the same individuals across multiple time points, is more informative but harder and more expensive to collect. We analyzed brain scans from over 6000 young people in the Adolescent Brain Cognitive Development (ABCD) study, about half of whom were girls. Each participant had brain scans at the start of the study, two and four years later. We showed that Baseline-Conditioned Norms (B-Norms), used each person’s first scan and their ages at baseline and follow-up timepoint to predict later brain changes. We compared this to Cross-Sectional Norms (C-Norm), which only used age. B-Norms predicted brain thickness more accurately and importantly were better at detecting brain differences linked to puberty, especially in girls. Although better fit alone does not prove superiority, our findings suggest that using our proposed B-Norms, we potentially also capture more developmental variance suggesting that our B-Norms are possibly more sensitive to sex-specific brain development over time.

## Background

The predominant analytical approach in neuroimaging for identifying disease-related brain alterations has been case-control comparisons. Over the past decade, this group-level approach has increasingly been complemented by normative modelling - a framework that enables the quantification of individual deviations from expected neurobiological patterns relative to a reference population (1,2). In brief, normative models estimate centiles of variation across neurobiological or behavioral measures, allowing inferences about an atypical development at the individual level without the need for predefined diagnostic groups.

A key strength of normative modelling lies in its flexibility to model diverse mappings across different phenotypic domains, ranging from neuroimaging measures (e.g., cortical thickness/volume) to behavioral or demographic traits (3). Therefore, this modeling approach is particularly well-suited for lifespan research, where inter-individual variability often reflects subtle and complex developmental or degenerative processes (3–6). For example, deviations from normative neurodevelopmental trajectories have been implicated in the pathogenesis of psychiatric and neurodevelopmental conditions (7). More recent applications have used normative models to investigate cognitive decline and deterioration or increases of structural brain measures (e.g., cortical thickness/volume) in aging and neurodegenerative disorders (4,8–13). These models further provided valuable insights into heterogeneity in psychiatric conditions such as attention-deficit/hyperactivity disorder (14,15), schizophrenia (14,16–18), autism spectrum disorder (16,19–22), and Alzheimer’s/dementia (23–27).

Despite these advances, most normative modelling studies have relied on cross-sectional data. To some extent, this limits their capacity to accurately predict within-subject longitudinal change. A recent study showed evidence that cross-sectional models may underestimate dynamic brain changes over time (8,28,29). Although recent studies have begun to incorporate and investigate longitudinal data (e.g., (30)), some of which, however, still apply cross-sectional models to individual data points, rather than leveraging longitudinal information (27,31,32). Among other disciplines developmental neuroscience has emphasized the need for longitudinal designs and investigations to capture changes during brain maturation - a crucial developmental period for which mounting evidence of vulnerability to psychiatric illness has been put forth (33–37). Recent findings suggest that modelling deviations in the timing of pubertal onset - such as early puberty onset - can enhance the prediction of later mental health outcomes (34,38). Such evidence highlights the translational potential of (longitudinal) normative models for early detection and thus possible interventions.

To this end, we estimated Baseline-Conditioned Norms (B-Norms) that use baseline cortical thickness data and age to predict cortical thickness at later timepoints and compared them to Cross-sectional Norms (C-Norms). Specifically, we used data from the Adolescent Brain Cognitive Development (ABCD) Study (39), comprising baseline data from 5374 participants (2515 female; baseline: mean±SD age = 118.78±7.44 months), and data at 2-year follow-up which were used to train our models. Furthermore, we used data from participants who participated at three timepoints to test our models: 805 participants (366 female; baseline: mean±SD age = 119.28±7.34; 2-year: mean±SD age = 143.22±7.49; 4-year: mean±SD age=168.45±7.77). Cortical thickness measurements were derived from 360 regions of interest (ROIs) based on the Glasser atlas (Glasser, 2016). For both B- and C-Norm models, we trained them for each brain region separately, hypothesizing that B-Norm models utilizing longitudinal data would show enhanced sensitivity to developmental changes within the respective developmental time period. To validate this, we examined the relationship between model-derived deviation scores and pubertal development as determined by the Pubertal Development Scale (PDS) scores. Validation was performed in the held-out test set. PDS scores acquired at 2-year and 4-year follow-up were statistically associated with the respective model-derived deviation scores of C- and B-Norm models. Our findings aim to contribute to the ongoing refinement of normative modelling approaches, especially in the context of individual-level longitudinal developmental predictions.

## Methods

### Dataset

We made use of longitudinally acquired data of the Adolescent Brain Cognitive Development (ABCD, Casey et al., 2018) study. In the ABCD dataset children are recruited at ages 9 to 10 with the aim of characterizing brain developmental trajectories. To this date more than 11.868 children were recruited across 21 different sites in the United States of America. Study procedures have been approved by either the local site Institutional Review Board (IRB) or by local IRB reliance agreements with the central IRB at the University of California. All participants and their parents or legal caregivers provided written informed consent. Data for the current study was obtained from ABCD release 5.1 utilizing phenotypic and imaging data from the baseline, 2-year, and 4-year follow-up visits.

### Data selection and preprocessing

#### Demographics

The ABCD project provides a multitude of tabulated data. Here, we made use of the following files: *abcd_p_demo* to extract sex and ethnicity; *abcd_y_lt* to extract the interview age at baseline and follow-up visits; the *ph_y_anthro* file to compute body mass index (BMI) which we use to exclude participants with unusually large BMI; and *mri_y_adm_info* for information about scan sites which we use as covariates during model training and prediction. We additionally computed mean puberty score (PDS) as rated by the youths’ caregivers from data of the *ph_p_pds* file to associate deviation scores with youth’s pubertal progress.

#### Puberty scores

We calculated a summary statistic representing progress in pubertal development using items of the Pubertal Development Scale (PDS; Herting et al., 2021; Kraft et al., 2023). This rating scale was designed to reflect Tanner stages without the need for physical examination (42,43). In this questionnaire, a child’s pubertal development is assessed using a four-point Likert scale ranging from ‘has not begun’ to ‘completed’. These items were specific to certain physical characteristics (including skin changes, breast development, deepening in voice, etc.). Please note that some items were administered based on sex. For example, the onset of menarche was exclusively asked for females and is a binary (i.e., either 1 or 4) rating. The ratings are either provided by the children or their caregivers. In this study, we focus on ratings provided by the caregivers for two reasons: a) the self-reported ratings appeared less reliable and b) more data is available for caregiver ratings (42).

#### Cortical thickness data

We preprocessed the raw structural data of the ABCD on an in-house cluster computer (Ubuntu 22.04) using the recon-all functionality of Freesurfer v7.4.1 (44). Cortical thickness values were calculated and extracted for 180 regions per hemisphere as defined by the Glasser atlas (45). In addition, we stored Euler numbers which we used to exclude badly reconstructed data during preprocessing.

#### Preprocessing

Before preprocessing, data comprised of 11868 participants with a baseline measure, 10908 participants with 2-year follow-up data, and 4688 participants with 4-year follow-up data. We excluded all those participants which had only a single measurement. Additionally, we excluded all participants with BMIs of less than 10 or larger than 50. We included this step because BMI has previously been associated with changes in cortical thickness in adolescents (46) as well as in adults (47,48). Furthermore, we excluded all those participants who had no valid or no pubertal development score (PDS) rating and cortical thickness measures available. We then separated those participants with only baseline- and 2-year follow-up data into a subset and those with baseline-, 2-year, and 4-year data into another. For both subsets we used the Euler numbers and computed their mean and standard deviation for each of the ABCDs scan-sites. We then excluded all participants whose Euler numbers were larger than 6 SDs from the mean for a given scan-site. This was to ensure that no extremely badly reconstructed data was used (for overview plots see Supplementary Figures S1 and S2). Lastly, we performed a similar exclusion step for cortical thickness values. We again calculated the mean cortical thickness and standard deviation across subjects for each ROI and excluded all participants who exceeded the 6 SDs threshold. This was done to prevent strange overfitting phenomena due to extreme outliers. For details on the stepwise number of excluded participants see **Figure 1** below.

**Fig 1.**
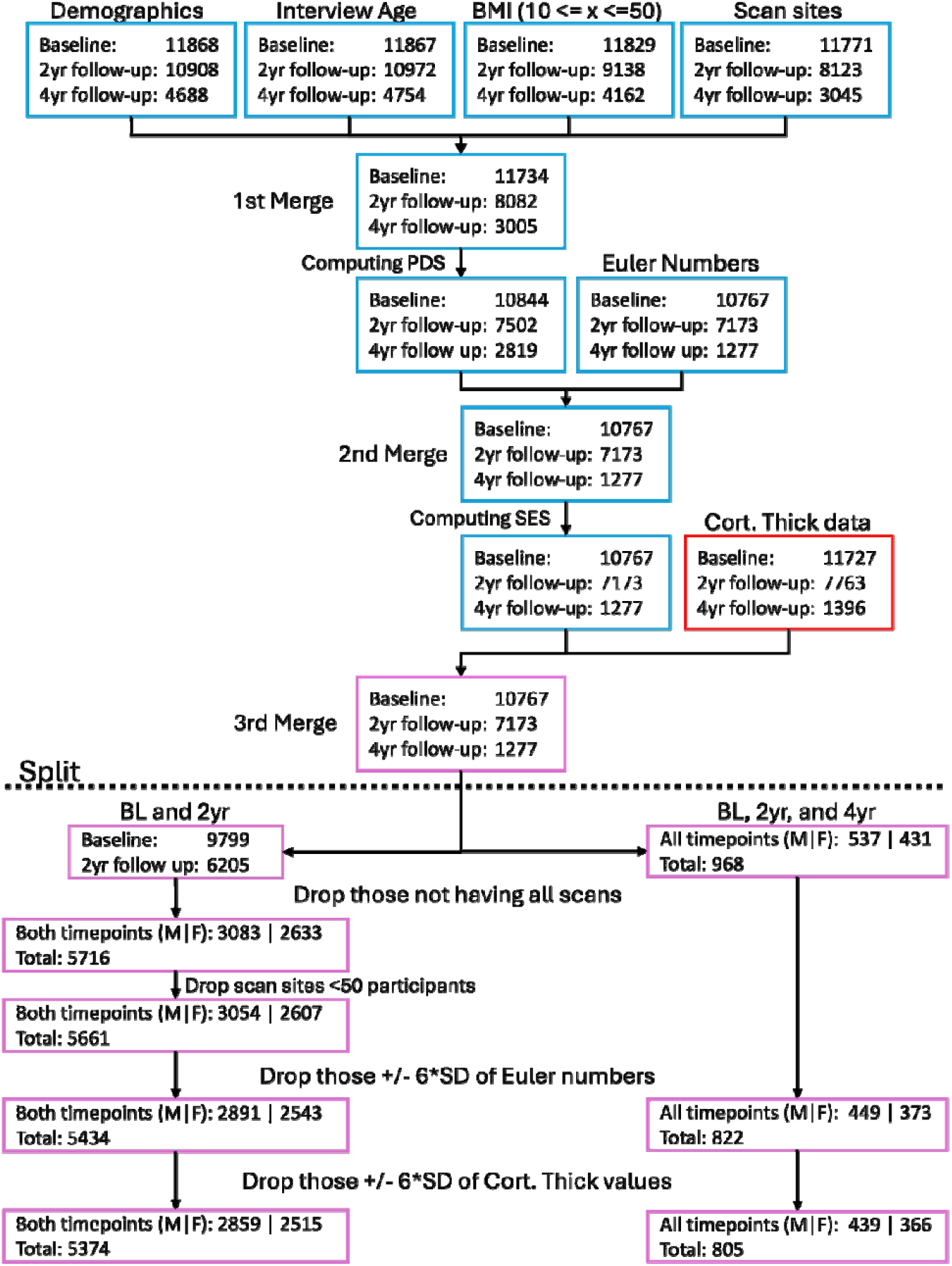
Data preprocessing flowchart of the ABCD dataset. We first loaded data from the ABCD study. Namely demographics (*abcd_p_demo*), interview age (*abcd_y_lt*), we computed BMI (*ph_y_anthro*) and excluded all participants that had a BMI between 10 and 50 and loaded the scan site information (*mri_y_adm_info*). We then merged these files (1^st^ Merge). After that, we computed PDS scores (*ph_p_pds*) as described previously (CITE). The resulting data was then merged with Euler Numbers (obtained from inhouse Freesurfer reconstruction; 2^nd^ Merge). After that we computed socioeconomic status (*abcd_p_demo*) and merged the resulting data with the cortical thickness data we received from our inhouse Freesurfer preprocessing. This resulted in the data represented by “3^rd^ Merge”. We then retained data for all participants who had Baseline, 2yr-, and 4-year follow-up data (right path below the dashed “Split” line). For the remaining data (left path below the “Split” line), we first retained only those participants who had both Baseline and 2yr follow-up data. For both paths, we then removed those participants whose Euler Numbers exceeded 6*SD from the mean. Lastly, we removed all those participants who had unusually large (again exceeding 6*SDs from the mean) cortical thickness values *within* any Glasser brain region.

##### Train test and splits

After preprocessing we designated those participants with only baseline (female: N=2515, age=118.30±7.39 [mean±SD]; male: N=2859, age=119.06±7.49) and 2-year follow-up measurements (female: age=142.92±7.81; male: age=143.74±7.81) as the training dataset. Participants with baseline (female: N=366, age=118.87±7.47; male: N=439, age=119.62±7.41), 2-year follow-up (female: age=142.73±7.31; male: age=143.64±7.62), and 4-year follow-up (female: age=167.82±7.48; male: age=168.98±7.99) data were used for testing. This approach ensured independent samples during the training process as well as for testing. The splits did not differ significantly on core demographic variables such as age or BMI at baseline but showed some statistical differences as determined by a Welch’s t-test in PDS scores. We argue that this difference, while unfortunate, is barely avoidable as stratifying for these variables would have resulted in a much smaller test set. For more details see supplementary figures S3-5 and the corresponding text.

### Normative modelling with Bayesian Linear Regression

We employed normative modeling using Python 3.12.9 and the PCNtoolkit (49) package (version 0.33). Bayesian Linear Regression (BLR) with likelihood warping (50) was used to predict cortical thickness from a covariance matrix including “age_bl_ and site” for the standard Cross-Sectional normative models (C-Norms) and “cortical thickness_bl_, age_bl_, age_follow-up_, and site” for the Baseline-Conditioned normative models (B-Norms). Input age values were the age in months. Sinarcsinsh was employed for warping (51). Site covariates were encoded as one-hot vectors (i.e., one vector per site where the respective column was set to “1” for a particular site). For each of the 180 brain regions for both hemispheres as defined by the Glasser atlas (45), cortical thickness is predicted as:

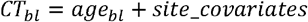

for the C-Norms, and

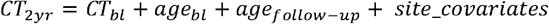

for the B-Norms. Where the subscript ‘*follow-up*’ in the second age-parameter corresponds to either the age at the 2-year or 4-year follow-up visit. While the age variables certainly are correlated to one other, we argue using them this way is not of a particular issue as we are not interpreting the contribution of individual variables

Both models were optimized using the powell algorithm, and results are based on models trained on the training split and evaluated on the independent test set. We assessed the model fit for each brain region using several metrics, including Pearson’s correlation between observed and predicted measures, root-mean-squared error (RMSE), standardized mean-squared error (SMSE), explained variance (EV), and mean squared log-loss (MSLL). Additionally, we evaluated skewness and kurtosis to estimate higher-order moments beyond the mean in the test set (52).

### Comparing performance measures

We used the ANOVA function from pingouin python package to perform 2-factor repeated-measures analysis of variance (rm-ANOVA) to statistically assess differences in performance metrics of the C- and B-Norms separately for both males and females. The dependent variables were the performances measures as defined earlier. Our within-subject factors were MODEL (levels: C-Norm and B-Norm) and TIMEPOINT (levels: 2-year and 4-year follow-up), the identifier variable were the regions of interest as defined by the Glasser atlas. We show statistics for explained variance, skewness, and kurtosis in the main text. Mean squared log loss (MSLL), (standardized) mean squared error ([S]MSE), root-mean squared error (RMSE), and Bayesian Information Criteria (BIC) are presented in the supplementary material.

### Validations of normative models against puberty scores

We performed an association analysis to reveal relationships between puberty scores as rated by the youths’ caregivers and the deviation scores (i.e., z-scores) obtained from the normative models. We conducted this analysis performing generalized linear models (GLMs) with a Gaussian family and identity link using the statsmodels Python-package, specifically the glm function. For each sex-specific model and each ROI, we computed the association *PDS*_*t*_ *~ zROI*_*t*_, where *PDS* are vectorized mean PDS scores per participants and *zROI* is a vector containing subjects’ deviation scores as obtained by the C- or B-Norms at follow-up time point *t* (i.e., 2-year or 4-year). This resulted in 360 z-statistics and p-values (which we corrected for multiple comparisons using the Benjamini-Hochberg (53) false discovery method), one for each ROI, for each model, and sex. Model degrees of freedom (df) were 1 and residual df differed between sexes (437 for males and 364 for females). Deviance and scale are reported in the results section. Statistics were then projected onto a surface brain and thresholded according to the critical BH values. As a supplementary analysis, we included BMI and socioeconomic status as covariates. These results are available in the supplementary materials. Furthermore, we provide the results in tabular format as supplements.

### Validation of normative models using puberty subgroups

To examine differences in positive and negative deviations across ROIs, participants were grouped into five pubertal stages (pre-, early-, mid-, late-, and post-pubertal), following Herting et al. and Kraft et al. (40,41). Glasser ROIs were aggregated into six lobe-level regions: occipital, frontal, temporal, parietal, insular, and cingulate – based on previously proposed definitions (54). For each participant and lobe, we counted the number of regions exceeding z-scores of +/−1.96, producing lobe-specific deviation vectors. These count vectors were stratified by pubertal stage, and Kruskal-Wallis tests were conducted across stages for each lobe, deviation type (positive/negative), model (cross-sectional/Baseline-Conditioned), and timepoint (2- and 4-year follow-ups), totaling 192 tests.

### Association of stable, positive or negative percentile shifts with Puberty progression

We additionally validated the normative models by categorizing participants into three groups based on percentile shifts between the 2-year and 4-year follow-up data. Specifically, we defined three groups: negative (zDiff < −1), stable (−1 < zDiff < 1), and positive (zDiff > 1), with zDiff representing the change in ROI-specific z-scores over time. For each ROI, participants were assigned to one of these groups; the number of participants per group therefore varies. For each ROI a Kruskal-Wallis test was conducted to assess group differences in delta PDS scores (4-year minus 2-year). Please note: the results of this analysis are available in Section 5 of the supplements.

## Results

### Integrating Baseline Thickness Measures as Predictors for Norms in Longitudinal Designs

While C-Norms predict cortical thickness based on age, we here examined if a different modelling approach by using cortical thickness at baseline alongside age at follow-up as features yields more accurate predictions (B-Norms), and if the resulting deviation scores yield stronger and potentially more meaningful associations with external variables that are sensitive to change, such as pubertal development. We trained B-Norms and C-Norm models in the same training data set, separately for males and females to account for potential sex specific trajectories. To validate the trained models, we compared explained variance for our models when applied to the 2-year and 4-year follow-up held-out test data. This allowed us to assess the accuracy of fits (i.e., the center of the distribution). Additionally, we assessed differences in higher order moments (i.e., skewness and kurtosis) of the resulting deviations (i.e., z-scores) to investigate the shape and calibration of the centiles of deviation. The latter is important as it ensures that the models accurately fit to the overall distribution and thereby allow for reliable inferences (55). While the comparison between the two models is not of most importance for this article, we include it here for completeness. We found marked differences in the mean fit between models. Given that we use baseline cortical thickness as a prediction parameter in our B-Norm models, we expected to find more explained variance for both 2-year and 4-year follow-up data compared to C-Norms, which indeed we did (**Figure 2A**). The overall pattern was similar across sexes, with higher variance explained for B-Norms (mean±SD across timepoints, males: 0.635±0.125; females: 0.633±0.123) compared to C-Norms (males: 0.023±0.038; females: 0.026±0.04). On top of these large B-vs-C-Norm differences, we observed a small model-by-timepoint interaction, indicating that the accuracy of mean fits decreased from 2-year to 4-year follow up, with largest effects in B-Norms and in males (model-by-timepoint interaction, males: F_1,359_ =25.86, p<5.91e^−7^, η^2^=0.0672; females: F_1,359_ =26.04, p<5.43e^−7^, η^2^=0.0676; post-hoc t-tests males: *C-Norm* mean-diff=0.0038, t=3.268, p<1.18e^−3^, *B-Norm* mean-diff=0.0154, t=7.016, p<1.14e^−11^; post-hoc tests in females: *C-Norm* mean-diff=0.0006, t=0.613, p<0.54; *B-Norm*: mean-diff=0.0129, t=5.618, p<3.88e^−8^).

**Fig 2.**
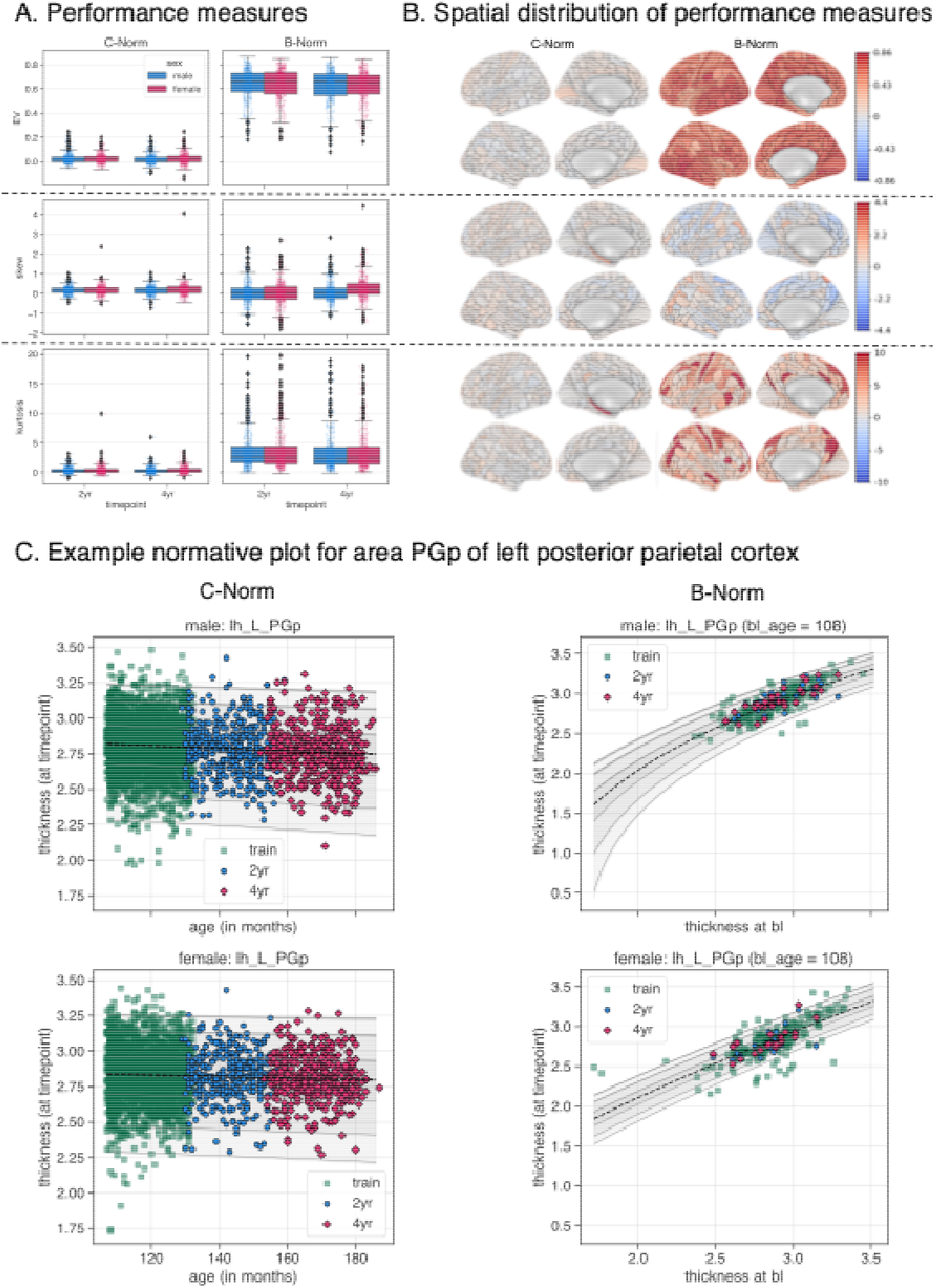
Cross Sectional Norms (C-Norm)-vs Baseline-Conditioned Norms (B-Norm). Panel A) depicts three performance measures of the different normative models: explained variance (top row), skew (middle row), and kurtosis (bottom row) for both models (columns). In panel B) the first two columns correspond to the lateral and medial views of the C-Norms, the last two to the B-Norms. Warmer colors indicate higher explained variance, positive skewness or kurtosis; colder colors indicate negative skewness or kurtosis. Panel C) shows normative plots of an example ROI (here the PGp, as defined by the Glasser atlas). The columns represent normative trajectories for the standard C-Norms (left) and the B-Norms (right). Rows correspond to sex. Within each graph, green squares indicate the training data. The black dashed lines indicate the median. The blue circles and red diamonds correspond to the 2-year and 4-year follow-up data, respectively. Grey lines indicate the centiles (1%, 5%, 25%, 75%, 95%, and 99%) and gray patches around these lines indicate the respective uncertainty. Note: given the nature of the B-Norms, we needed to plot one normative plot per baseline age. Only one baseline age (108 months) is depicted here. Others are available in the supplementary material (see Figure S20).

For the shape of the distribution and the centiles of the predicted z-scores, we investigated skewness and excess kurtosis (i.e., kurtosis values below or greater than 0), where values close to zero would indicate a standard normal distribution. On average, we found that skewness was closer to zero for B-Norms than C-Norms, however, variance was larger in B-Norms. We observed larger excess kurtosis for B-Norms as compared to C-Norms. The subpanels in **Figure 2A** depict the distributions and the corresponding statistics for explained variance, skewness, and kurtosis, among other statistics, from our repeated measures ANOVA are detailed in the Supplement (see Figures S6-19 and Tables S1-8). **Figure 2B** illustrates the spatial distribution of the performance metrics, exemplarily for females at 2-year follow-up. In the depicted female B-Norms, we found highest explained variance in large parts of the occipital (left/right PIT, VMV1, and left MT) and temporal regions (right PHT, PH, TP0J2 and left TP0J1, TE1m and STSda), and lowest explained variance scores in frontal (left/right pOFC, right OFC, 13l, and 25 and left 6d) and insular (left AAIC and PoI2, and left/right FOP3) areas. Similar results were found for the female 4-year follow-up data. Results for male B-Norms, were mostly consistent with females (see supplementary Figures 21-22). Interestingly male C-Norms showed different highest explained variance scores in parietal and frontal areas for the 2-year and 4-year follow-up data. Furthermore, we saw differences between lowest explained variances in male C-Norms for the 2-year and 4-year follow-up data. Specifically, we found different areas of the frontal lobe that showed explained variances below 0 (i.e., the model performs worse than chance) for the 2- and 4-year follow-up data as well as some temporal regions in the 2-year and parietal regions in the 4-year follow-up data. These results were inconsistent with the predictions made by female C-Norms on 2-year follow-up data with largest explained variance in frontal and temporal regions. Supplementary Figures S21 and 22 provide surface maps for the female 4-year and the male 2- and 4-year data. Supplementary file *test_metrics*.*csv* provides all metrics. Largest and smallest skewness and excess kurtosis also differed between male and female models for both timepoints. These results suggest that there may be sex differences in estimating ROI-wise C- or B-Norms.

ROI-wise comparisons of mean fits for the C-Norms and B-Norms reveal differential effects, that is, some ROI models explained more variance in the 2-year than in the 4-year follow-up or vice versa. For example, in the female longitudinal B-Norm model for a section of the posterior cingulate cortex (i.e., 31pv) of the right hemisphere, we found higher explained variance in the 2-year than in the 4-year follow-up data, whereas for the right PEF (premotor eye field) ROI the model explained more variance in the 4-year follow-up data. The full range of these results is available in the supplements (see supplementary Figures S8, 9, 14, 15, 17, and 18).

### Association with puberty scores

We investigated whether deviation scores of the 2-year or 4-year follow-up test data were associated with puberty stage as rated by the youths’ caregivers. Specifically, for each of the 360 ROIs, for each model and for each sex, we tested for linear associations, using generalized linear models (GLMs), between pubertal scores (PDS) and the deviations (z-scores) from B-Norm and C-Norm model, with Benjamini-Hochberg (BH) false discovery correction applied across tests.

For C-Norms, only one association survived correction for multiple comparisons; specifically in 2-year follow-up data for females in the left hemisphere’s area 31pv, a part of the left posterior cingulate cortex (z=-3.985, p<0.02442, BH corrected, N=366, df_model_=1, df_resid_=364, scale=0.41, deviance=149.5). For female B-Norms, four areas of the left hemisphere survived correction; specifically for two subsections of the lateral occipital cortex (LO2: z=-3.597, p<0.0362, BH corrected, df_model_=1, df_resid_=364, scale=0.168, deviance=61.15 and LO3: z=-3.540, p<0.0362, BH corrected, df_model_=1, df_resid_=364, scale=0.1682, deviance=61.22), an area in the lateral frontal lobe (IFSa: z=-3.619, p<0.0362, BH corrected, df_model_=1, df_resid_=364, scale=0.168, deviance=61.13), and an area of the insula (PoI2: z=3.632, p<0.0362, BH corrected, df_model_=1, df_resid_=364, scale=0.168, deviance=61.11). These results suggest, that with progression through puberty, areas LO2, LO3, and the IFSa exhibit decrease in cortical thickness, whereas parts of the insula appeared to show positive deviation. A corresponding surface map for the significant B-Norm areas is depicted in **Figure 3A** (for the 31pv of C-Norms see supplementary figure S23). Furthermore, the distributions of t-scores across ROIs in **Figure 3B** suggest that there is an average negative shift for the females in the 2-year data for both the C-Norms and B-Norms. This negative shift for the females is lost for the C-Norms in the 4-year follow-up data but remains for the B-Norms, indicating that such models may better capture a relationship between puberty progression and changes in cortical thickness.

**Fig 3.**
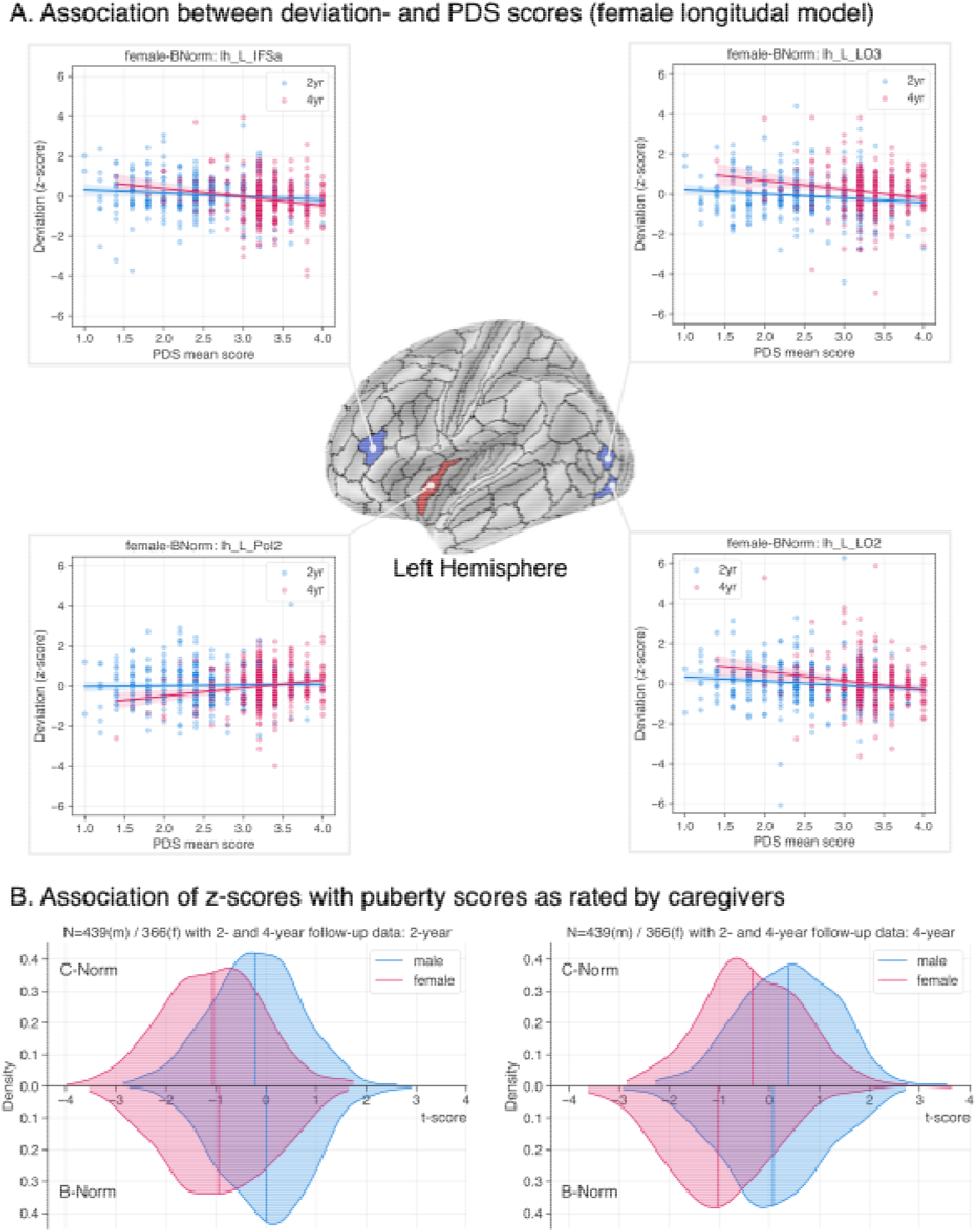
Association of deviation scores with puberty. Panel A shows a spatial surface map with the statistically significant areas for the female Baseline-Conditioned normative (B-Norm) models and the respective scatter plots and regression lines. The three areas marked in blue correspond to negative z-statistics as obtained from a GLM testing “PDS ~ z”, whereas the red marked area corresponds to the positive z-score. Blue (2-year) and red (4-year) colored dots within the scatter plots correspond to a single individual. Colored lines reflect the respective regression line of the data; patches around lines indicate the 95th confidence interval. The scatter plot for the significant female cross-sectional normative (C-Norm) model is available in the supplementary material (Figure S23). Panel B shows the overall distributions of the GLM z-statistics for this analysis. Colors indicate sex (red = female, blue = male), curves above the x-axis correspond to the associations for the C-Norms, whereas those below correspond to B-Norms.

We also performed the same analysis using the simple difference between the 2- or 4-year data and baseline cortical thickness (i.e., 2-/4-year minus baseline). This is most common approach for longitudinal data but yielded no significant associations. The ROI-wise statistics for this approach are available as supplementary csv files.

In addition, we adjusted our train- and test set for family structure since the ABCD dataset has high proportions of siblings/twins. For this approach, we found no difference in performance and predicted norms between the models training using all subjects and those trained with data accounted for family structure (see Supplements section 4.1.). However, we did not find the previously reported statistically significant associations in the aforementioned brain areas. Significance values increased to p=0.0965 for the left 31pv and to p=0.0636 for the left LO2, IFSa, PoI2 and LO3. This could be because sample sizes decreased by 37 and 26 in the males (new N=402) and females (new N=340) respectively.

Lastly, we added BMI and socioeconomic status (SES) as covariates in our GLM analyses. That is, the formula for the GLM then was “PDS~z*BMI*SES” yielding main as well as interaction effects. Including these variables no longer yields significant associations between deviation and PDS scores. However), BMI was highly associated with PDS in nearly all brain regions for both males and females (56) as well as both follow-up timepoints, which poses issues regarding co-linearity. SES was also associated in nearly all brain areas, however only for both male and female CNorm-2year data, for female-Cnorm-2year data, and for both male and female BNorm-2year data. Full tabular statistical data files for these analyses are also available as supplementary files.

### Validation of normative models using puberty subgroups

We further evaluated whether positive or negative deviation counts, defined as the sum of regions with deviation scores greater than or less than z=±1.96, were associated with specific pubertal stages. To this end, we grouped participants according to their pubertal stage (pre, early, mid, late and post pubertal, **Figure 4A**) and analyzed deviations at the level of lobes (**Figure 4B**). For females (see **Figure 4C** top), 11 Kruskal-Wallis omnibus tests yielded significant group effects. In cross-sectional normative models (C-Norms), marginally significant group differences were found among negative deviators at the 2-year follow-up in the left occipital (H(4)=9.96, p=0.041) and parietal (H(4)=14.38, p=0.0062) lobes; Dunn’s post-hoc tests suggested a mid-vs late-pubertal group difference in the parietal lobe (p=0.0053) but not in any of the pubertal stages for the occipital lobe (min p=0.139 for mid-vs late-pubertal). For positive deviators, we found significant effects in the left cingulate for the 2-year follow-up data (H(4)=15.33, p=0.0041), with early-vs late-(p=0.036) and mid-vs late-pubertal (p=0.0101) contrasts, and in the right cingulate for the 4-year follow-up (H(2)=7.10, p=0.0288), though pairwise comparisons were not significant (min p=0.0705 for late-vs post-pubertal).

**Fig 4.**
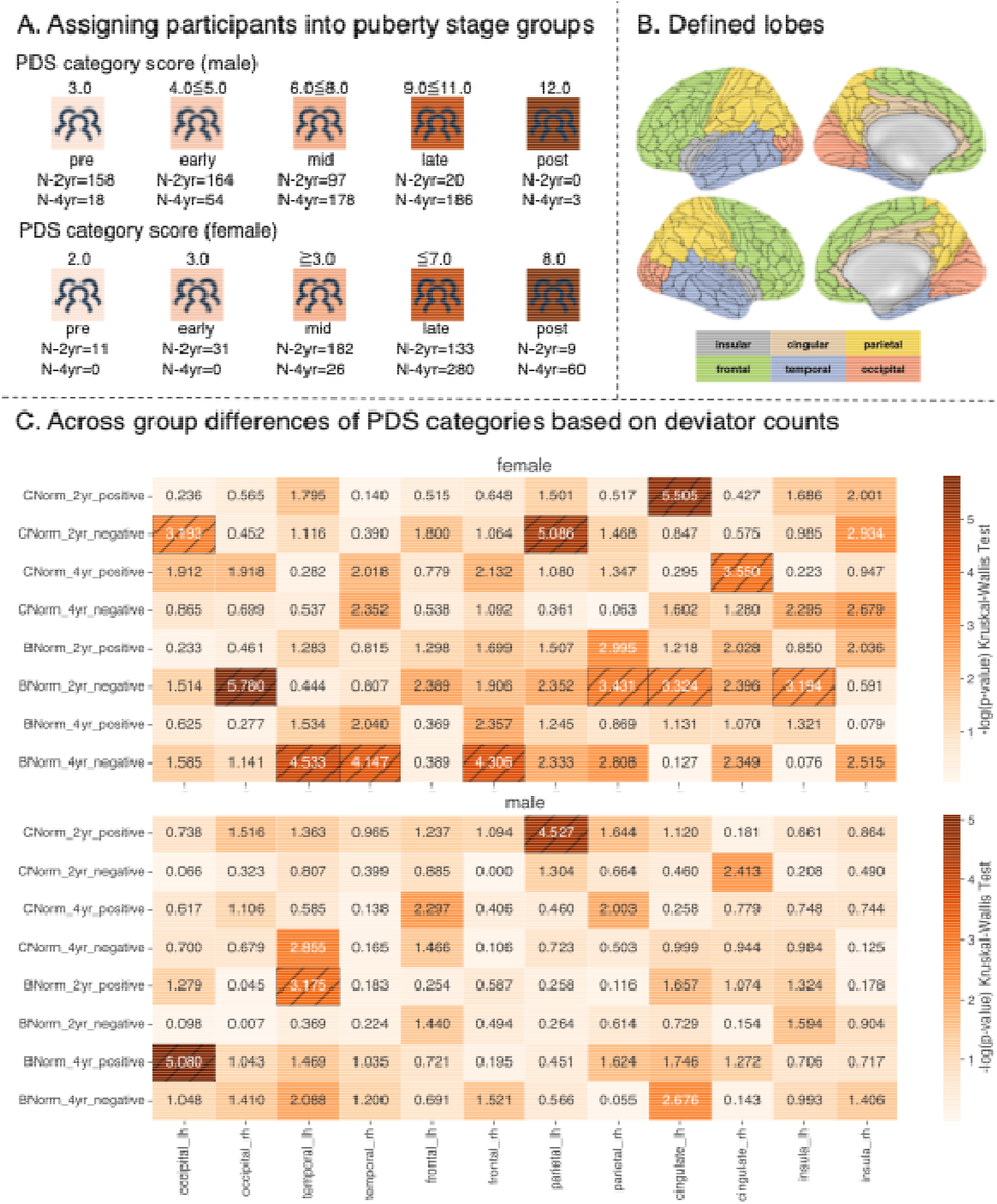
Definition and differences in pubertal stages for lobe-wise positive and negative deviation counts. Panel A) shows the definiton of pubertal stages/categories based on puberty scores (see Herting et al., and Kraft et al.). Panel B) illustrates which regions of the Glasser atlas were combined into lobar regions (definition based on (54). Panel C) shows −log-transformed p-values according to Kruskal-Wallis omnibus test for differences in positive/negative deviation counts between pubertal categories (pre-, early-, mid-, late-, and post-pubertal) for both timepoints using the C-Norms or B-Norms for female (top) and male (bottom) data. Hatched cells indicate FDR corrected significance.

B-Norms showed significant effects for negative deviators in the right occipital (H(4)=15.95, p=0.0031; mid-vs late-pubertal p=0.0031), right parietal (H(4)=10.53, p=0.0324; min pairwise p=0.069 for early-vs mid-pubertal), and the left cingulate (H(4)=10.276, p<0.0360, min p=0.221 for mid-vs late-pubertal) after two years. At 4-years follow-up, significant overall pubertal stage differences were found for negative deviators in the left (H(2)=9.07, p=0.0108) and right (H(2)=8.30, p=0.016) temporal lobes, and the right frontal lobe (H(2)=8.61, p=0.0135), all driven by mid-vs late-pubertal contrasts (p=0.095, p=0.012, and p=0.01, respectively). Corresponding p-values are visualized in **Figure 4C**; significant cells are hatched in black. We used Bonferroni correction to adjust p-values for multiple comparisons for Dunn’s post-hoc tests. These results suggest, that for females, the largest differences between pubertal stages in relation to extreme deviations from the norm can be found between mid- and late pubertal participants.

For males (see **Figure 4C** bottom), we only found three significant Kruskal-Wallis omnibus tests. That is, for the C-Norms, we found significant group differences only the left parietal lobe of negative deviators in the 2-year follow-up data (H(3)=11.176, p<0.0109, specifically for early-vs mid-pubertal p=0.0134). For B-Norms, we found marginally significant effects for positive deviators in the left temporal lobe (H(3)=8.213, p<0.042, specifically for pre-vs early pubertal p=0.0481) in the 2-year follow-up data and slightly stronger results in the left occipital lobe (H(4)=14.364, p<0.0062, specifically for mid-vs late-pubertal p=0.038) for 4-year follow-up data.

## Discussion

Our study shows that Baseline-Conditioned Norms (B-Norms), which model future cortical thickness measures based on baseline cortical thickness, baseline age, and age at follow-up, appeared to capture more meaningful associations between brain structure and pubertal development than Cross-Sectional Norms (C-Norms) in a longitudinal context such as the ABCD study (39). B-Norms revealed marginally significant region-specific deviation patterns associated with pubertal stage progression in females, particularly at later follow-ups, including four years after baseline. In contrast, these associations were absent or markedly weaker when using C-Norms. Despite being marginal effects, these results hint toward a benefit by priming models using baseline measures. Our perceived enhanced sensitivity of B-Norms likely stems from their ability to capture developmental changes occurring between baseline and follow-up, which are obscured when relying solely on age-based cross-sectional norms. Notably, the link between the number of negatively deviating regions within lobes and pubertal stage progression hints towards the potential of B-Norms to detect subtle, regionally specific brain changes during adolescence. However, these results need to be investigated further with more available data, e.g., using the next batch of the ABCD dataset.

Conversely, growth chart-style analyses did not reveal significant associations with pubertal status at this stage, suggesting that models using baseline cortical thickness measures to predict future thickness measures are potentially better suited for detecting nuanced developmental trajectories. Performance differences between the two model types indicated that B-Norms substantially outperformed C-Norms, explaining approximately 60% more variance in cortical thickness at both 2-year and 4-year follow-ups. However, given the design of our B-Norm model by using baseline measures of the variable of interest (i.e., cortical thickness), such performance differences are to be expected. It should be further noted that C-Norms in our analysis performed worse than what has been reported in prior work using similar models trained across the full lifespan (3). It is likely that the narrow age range of the ABCD study sample accounts for the lower performance overall. However, this age range attributed performance drop cannot explain the superiority of B-Norms over C-Norms in our work, given that the training data used for C-Norms and B-Norms is identical. Investigating the spatial distribution of performances across the brain surface allowed us to further investigate performance differences between C-Norms and B-Norms. While B-Norms achieved largest explained variances in the occipital (Glasser: left/right PIT, VMV1, and left MT) and temporal (Glasser: right PHT, PH, TP0J2 and left TP0J1, TE1m and STSda) regions with similar patterns across sexes, explained variance was lowest in parietal, insular and to some degree in frontal regions. Prior work linked parts of the occipital cortex to pubertal changes, including negative associations with testosterone in females (57). Although a positive association was reported in males, the relatively early pubertal stage of ABCD male participants may explain the absence of this effect here. Interestingly, Wierenga et al. (2022) reported an interaction of cortical thickness with age in the left insula for males (58). Here, our B-Norms fit insular regions less well in females, possibly reflecting more variability in females. Furthermore, we found inconsistent results in frontal and parietal regions for male and female C-Norms, areas previously tied to pubertal development (59). This could be a potential reason for the lower predictive performances of C-Norms compared to B-Norms.

As was highlighted in a recent commentary, it is important to investigate higher-order moments of model derived deviation score distributions in addition to the mean fit to properly assess model performance (55). Therefore, we also investigated differences in skewness and excess kurtosis for our C- and B-Norms. This analysis suggested that deviation scores obtained from C-Norm models were more normally distributed than for B-Norm models (excess kurtosis and skewness are closer to 0). The kurtosis results suggest that regional B-Norms are generally more sensitive to changes between visits, which can be seen by the narrower centiles of the normative plots as opposed to those of the C-Norms (see **Figure 2C**).

While better performance in some circumstances is desirable it is important to validate models using meaningful data. To this end, we examined whether deviation scores obtained from our models show a relationship with progression through puberty. The three regions that we found to be significantly associated all showed a negative association between pubertal process and deviation scores and were all in B-Norms of females. These negative associations with PDS may be consistent with previous findings in relation with estradiol levels (60). Area left PoI2 in the insula was associated with positive deviations (i.e., an increase in cortical thickness) with progression through puberty (higher PDS). Overall, this analysis suggested that deviation scores as obtained by our B-Norms were slightly more negatively associated with pubertal development in females at the 4-year timepoint than the cross-sectional models. This can be seen by the slight leftward shift of the respective distribution as seen on the right side of **Figure 3B**. These results suggest that while a better mean fit of the respective population technically means less variance in the residuals which often leads to fewer associations with phenotypic variables and may therefore not be useful in clinical settings. This issue has previously been reported for brain age prediction where models with very high fit can yield weaker associations with clinical variables (Bashyam et al., 2020 and see Hahn et al., 2021). Despite this effect our B-Norms still pick up on unexpected or unusual (e.g., early or late puberty onset in relation to the baseline cortical thickness and age) changes between the baseline and 4-year follow-up. Additionally, such changes could be quite small and yet lead to large deviations. This may be due to the high excess kurtosis, which indicates narrow distributions where even small changes produce large deviation scores. That these small changes align with pubertal progress suggests that incorporating baseline measures (B-Norms) increases sensitivity. However, given the small sample size of our test set these results should be interpreted with caution at this stage. Applying our models to the next ABCD release could shed more light on them. This would be interesting as longer intervals between baseline and follow-up may amplify this effect by detecting more unusual changes, which should be tested in future. In an auxiliary analysis we investigated whether family structure had an impact on this association analysis. After adjusting for family relationships in the training and testing set, the statistically significant associations mentioned before vanished. We argue that this could be due to the sample size drop in the test set since the performance and predicted norms of the models trained and evaluated using family-adjusted data did not differ from those obtained from models trained on all participants (see Supplements section 4).

Similar to previous studies that applied normative modelling to clinical populations, we investigated whether aggregated deviations from normative brain development, summed across regions of interest (similar to total outlier count, see Verdi et al., 2024; Wolfers et al., 2018), were sensitive to distinct pubertal stages. We observed sex-specific differences in how deviation counts varied across pubertal stages. Female participants showed greater effects: 4 lobes exhibited significant differences across stages in the C-Norms, and 7 in the B-Norms, compared to only 1 and 2 lobes respectively in males. Pairwise comparisons in the female 2-year follow-up cohort revealed that deviation scores differed significantly between mid- and late-pubertal stages in the right occipital and left cingulate lobes, and between early- and mid-pubertal stages in the right parietal lobe. Interestingly, in the 4-year follow-up data, negative deviations, which reflect cortical thinning relative to normative expectations, were observed in the bilateral temporal and right frontal lobes, primarily distinguishing mid-from late-pubertal females. These lobes are known to undergo significant cortical thinning during adolescence (63,64). Thus, our findings are in line with previously established findings that this process is stage specific, with late pubertal stages marked by increased deviation from normative developmental trajectories given a certain baseline and age. These findings align with previous literature emphasizing more dynamic neurodevelopmental trajectories in females during puberty, possibly due to earlier onset and more rapid progression of hormonal changes.

Reduced cortical thickness during puberty has been previously linked to synaptic pruning, myelination, and hormonal influences (65,66). In particular, thinning in temporal and frontal areas has been associated with cognitive maturation and may reflect the refinement of higher-order processes such as emotion regulation and social cognition. Such cognitive functions are especially sensitive to pubertal timing (67,68).

In a final validation step, using our normative models related to growth charts used in pediatrics, we categorized participants according to percentile shifts into negative, stable and positive deviators by computing the difference between the deviation scores (i.e., z-scores) between the 2- and 4-year follow-up data. This analysis did not reveal any significant effects after correcting for multiple comparisons rendering these results difficult to interpret (see Supplements section 5). We believe that one reason may be our chosen zDiff threshold – which is one of many possible thresholds – and the still relatively small sample size of our test set. Without correcting for multiple comparisons, we found marginal differences in associated regions between females and males (see Figures S25-26). Interestingly, if these results were to hold, ROI-wise male C-Norm and B-Norm models could show stronger and more differences than female models in this percentile shift analysis. This direction should be investigated further with future ABCD releases.

## Limitations

A limitation of this study, much like for any other current development of longitudinal methods, is restricted by the availability of large-scale datasets covering the age-range of interest. The degree to which our results translate beyond the population characteristics of the ABCD study cohort (39) remains to be investigated with future releases of new longitudinal datasets. Fortunately, such models can easily be extended, adapted, or retrained with new data releases.

Sample characteristics also apply when interpreting our results on sex differences. The majority of the male participants of the ABCD dataset did not yet fully progress through puberty, which likely explains the lack of puberty related associations with deviation scores. We argue that this has no implications for the results of the female models, as we have trained sex specific models and because it has been shown previously that brain development during puberty is different for males and females (58). A further limitation is that adding BMI and SES as confounds for our statistical analyses yields non-significant results. However, we are not the first to report a loss of effects incorporating these variables (41).

It should also be considered that at this point we cannot for certain attribute the loss of significant results in the association analysis when we accounted for familial relationship to either family-relationships or the drop in sample size. This should be investigated with future ABCD releases. Another limitation of the current study is the sole focus on puberty related changes. While interesting in their own rights, validation of C-& B-Norms using clinical variables (e.g., depression rating scales or similar) is necessary to estimate their utility in, for example, detecting and predicting developmental or age-related disease trajectories. Thus, future studies should incorporate clinical values and extended our proposed models. Furthermore, interpretation of the tested B-Norm models may be difficult due to their complexity. Given the required baseline features, cortical thickness and age as well as the age at a future timepoint, the model trajectories and centiles are estimated based on variable baseline cortical thickness depending on the baseline and future age. In theory this means that for each baseline and follow-up age per region of interest a growth chart must be generated (see Supplementary Figure S20 for two additional examples).

## Conclusion

In this study, we present a developmental application of normative modeling and demonstrate that Baseline-Conditioned Norms (B-Norms) utilizing longitudinal data yield deviation scores that are meaningfully associated with pubertal development. These associations were more pronounced in females than in males, likely reflecting the greater puberty-related variance in females in the respective age period. While still at an early stage, this Baseline-Conditioned modeling framework substantially improved model fit compared to cross-sectional approaches. These findings underscore the potential of this modeling approach to capture individual variability in different developmental phases.

## Supporting information

Additional statistics for all-subjects data

Additional statistics for family-adjusted data

## Ethics approval and consent to participate

Study procedures have been approved by either the local site Institutional Review Board (IRB) or by local IRB reliance agreements with the central IRB at the University of California. All participants and their parents or legal caregivers provided written informed consent.

## Acknowledgements

The study was funded by the Federal Ministry of Education and Research (Bundesministerium für Bildung und Forschung [BMBF]) and the ministry of Baden-Würtemberg within the initial phase of the German Center for Mental Health (DZPG) (grant: 01EE2306). T.K. and T.W. are members of the Machine Learning Cluster of Excellence, EXC number 2064/1 – Project number 39072764. TW acknowledges funding from German Research Foundation (DFG) Emmy Noether: 513851350. This work was supported by the BMBF-funded de.NBI Cloud within the German Network for Bioinformatics Infrastructure (de.NBI) (031A537B, 031A533A, 031A538A, 031A533B, 031A535A, 031A537C, 031A534A, 031A532B). The authors used data from the Adolescent Brain Cognitive Development^SM^ Study (ABCD, abcdstudy.org). ABCD data, held in the NIMH Data Archive (NDA), is a multisite, longitudinal study designed to recruit more than 10,000 children age 9–10 and follow them over 10 years into early adulthood. The ABCD Study® (https://abcdstudy.org/) is supported by the National Institutes of Health and additional federal partners under award numbers U01DA041048, U01DA050989, U01DA051016, U01DA041022, U01DA051018, U01DA051037, U01DA050987, U01DA041174, U01DA041106, U01DA041117, U01DA041028, U01DA041134, U01DA050988, U01DA051039, U01DA041156, U01DA041025, U01DA041120, U01DA051038, U01DA041148, U01DA041093, U01DA041089, U24DA041123, U24DA041147. A full list of supporters is available at https://abcdstudy.org/federal-partners.html. A listing of participating sites and a complete listing of the study investigators can be found at https://abcdstudy.org/consortium_members/. The ABCD consortium investigators designed and implemented the respective studies and/or provided data but did not participate in the analysis or writing of this report. This manuscript reflects the views of the authors and does not necessarily reflect the opinions or views of any other agency, organization, employer or company.

## Author Contributions

P.S. was responsible for data curation, code, analyses, and first draft of the manuscript. T.K. was responsible for preprocessing. T.K. and T.W. were responsible for conceptualization. All authors were responsible for writing and revising the manuscript.

## Availability of data and materials

Data from the Adolescent Brain Cognitive Development^SM^ Study (ABCD, abcdstudy.org) is available upon application on the NIH website.

Code is available on P.S. github-repository: https://github.com/PhilippS893/b-norm_modelling.

## References

1. Marquand AF, Rezek I, Buitelaar J, Beckmann CF. Understanding Heterogeneity in Clinical Cohorts Using Normative Models: Beyond Case-Control Studies. Biol Psychiatry. 2016 Oct;80(7):552–61.

2. Marquand AF, Kia SM, Zabihi M, Wolfers T, Buitelaar JK, Beckmann CF. Conceptualizing mental disorders as deviations from normative functioning. Mol Psychiatry. 2019 Oct;24(10):1415–24.

3. Rutherford S, Fraza C, Dinga R, Kia SM, Wolfers T, Zabihi M, et al. Charting brain growth and aging at high spatial precision. eLife. 2022 Feb 1;11:e72904.

4. Bethlehem RAI, Seidlitz J, White SR, Vogel JW, Anderson KM, Adamson C, et al. Brain charts for the human lifespan. Nature. 2022 Apr 21;604(7906):525–33.

5. Franke K, Gaser C. Ten Years of BrainAGE as a Neuroimaging Biomarker of Brain Aging: What Insights Have We Gained? Front Neurol [Internet]. 2019 [cited 2025 June 10];10. Available from: https://www.readcube.com/articles/10.3389%2Ffneur.2019.00789

6. Rutherford S, Barkema P, Tso IF, Sripada C, Beckmann CF, Ruhe HG, et al. Evidence for embracing normative modeling. eLife. 2023 Mar 13;12:e85082.

7. Insel TR. Mental Disorders in Childhood: Shifting the Focus From Behavioral Symptoms to Neurodevelopmental Trajectories. JAMA. 2014 May 7;311(17):1727.

8. Di Biase MA, Tian YE, Bethlehem RAI, Seidlitz J, Alexander-Bloch AaronF, Yeo BTT, et al. Mapping human brain charts cross-sectionally and longitudinally. Proc Natl Acad Sci. 2023 May 16;120(20):e2216798120.

9. Frangou S, Modabbernia A, Williams SCR, Papachristou E, Doucet GE, Agartz I, et al. Cortical thickness across the lifespan: Data from 17,075 healthy individuals aged 3–90 years. Hum Brain Mapp. 2022 Jan;43(1):431–51.

10. Jack CR, Knopman DS, Jagust WJ, Shaw LM, Aisen PS, Weiner MW, et al. Hypothetical model of dynamic biomarkers of the Alzheimer’s pathological cascade. Lancet Neurol. 2010 Jan;9(1):119–28.

11. Karas GB, Scheltens P, Rombouts SARB, Visser PJ, Van Schijndel RA, Fox NC, et al. Global and local gray matter loss in mild cognitive impairment and Alzheimer’s disease. NeuroImage. 2004 Oct;23(2):708–16.

12. Kjelkenes R, Wolfers T, Alnæs D, Norbom LB, Voldsbekk I, Holm M, et al. Deviations from normative brain white and gray matter structure are associated with psychopathology in youth. Dev Cogn Neurosci. 2022 Dec;58:101173.

13. Kjelkenes R, Wolfers T, Alnæs D, Van Der Meer D, Pedersen ML, Dahl A, et al. Mapping Normative Trajectories of Cognitive Function and Its Relation to Psychopathology Symptoms and Genetic Risk in Youth. Biol Psychiatry Glob Open Sci. 2023 Apr;3(2):255–63.

14. Kia SM, Marquand AF. Neural Processes Mixed-Effect Models for Deep Normative Modeling of Clinical Neuroimaging Data [Internet]. arXiv; 2019 [cited 2025 May 13]. Available from: http://arxiv.org/abs/1812.04998

15. Wolfers T, Beckmann CF, Hoogman M, Buitelaar JK, Franke B, Marquand AF. Individual differences v. the average patient: mapping the heterogeneity in ADHD using normative models. Psychol Med. 2020 Jan;50(2):314–23.

16. Pinaya WHL, Mechelli A, Sato JR. Using deep autoencoders to identify abnormal brain structural patterns in neuropsychiatric disorders: A large-scale multi-sample study. Hum Brain Mapp. 2019;40(3):944–54.

17. Wolfers T, Doan NT, Kaufmann T, Alnæs D, Moberget T, Agartz I, et al. Mapping the Heterogeneous Phenotype of Schizophrenia and Bipolar Disorder Using Normative Models. JAMA Psychiatry. 2018 Nov 1;75(11):1146.

18. Wolfers T, Rokicki J, Alnæs D, Berthet P, Agartz I, Kia SM, et al. Replicating extensive brain structural heterogeneity in individuals with schizophrenia and bipolar disorder. Hum Brain Mapp. 2021;42(8):2546–55.

19. Bethlehem RAI, Seidlitz J, Romero-Garcia R, Trakoshis S, Dumas G, Lombardo MV. A normative modelling approach reveals age-atypical cortical thickness in a subgroup of males with autism spectrum disorder. Commun Biol. 2020 Sept 4;3(1):486.

20. Wolfers T, Floris DL, Dinga R, Van Rooij D, Isakoglou C, Kia SM, et al. From pattern classification to stratification: towards conceptualizing the heterogeneity of Autism Spectrum Disorder. Neurosci Biobehav Rev. 2019 Sept;104:240–54.

21. Zabihi M, Oldehinkel M, Wolfers T, Frouin V, Goyard D, Loth E, et al. Dissecting the Heterogeneous Cortical Anatomy of Autism Spectrum Disorder Using Normative Models. Biol Psychiatry Cogn Neurosci Neuroimaging. 2019 June;4(6):567–78.

22. Zabihi M, Floris DL, Kia SM, Wolfers T, Tillmann J, Arenas AL, et al. Fractionating autism based on neuroanatomical normative modeling. Transl Psychiatry. 2020 Nov 6;10(1):384.

23. Alden EC, Lundt ES, Twohy EL, Christianson TJ, Kremers WK, Machulda MM, et al. Mayo normative studies: A conditional normative model for longitudinal change on the Auditory Verbal Learning Test and preliminary validation in preclinical Alzheimer’s disease. Alzheimers Dement Diagn Assess Dis Monit. 2022 Jan;14(1):e12325.

24. Pinaya WHL, Scarpazza C, Garcia-Dias R, Vieira S, Baecker L, F Da Costa P, et al. Using normative modelling to detect disease progression in mild cognitive impairment and Alzheimer’s disease in a cross-sectional multi-cohort study. Sci Rep. 2021 Aug 3;11(1):15746.

25. Verdi S, Marquand AF, Schott JM, Cole JH. Beyond the average patient: how neuroimaging models can address heterogeneity in dementia. Brain. 2021 Nov 29;144(10):2946–53.

26. Verdi S, Kia SM, Yong KXX, Tosun D, Schott JM, Marquand AF, et al. Revealing Individual Neuroanatomical Heterogeneity in Alzheimer Disease Using Neuroanatomical Normative Modeling. Neurology [Internet]. 2023 June 13 [cited 2025 Mar 10];100(24). Available from: https://www.neurology.org/doi/10.1212/WNL.0000000000207298

27. Verdi S, Rutherford S, Fraza C, Tosun D, Altmann A, Raket LL, et al. Personalizing progressive changes to brain structure in Alzheimer’s disease using normative modeling. Alzheimers Dement. 2024 Oct;20(10):6998–7012.

28. Korbmacher M, Vidal-Pineiro D, Wang MY, Van Der Meer D, Wolfers T, Nakua H, et al. Cross-sectional brain age assessments are limited in predicting future brain change [Internet]. Neuroscience; 2024 [cited 2025 June 10]. Available from: http://biorxiv.org/lookup/doi/10.1101/2024.09.11.612523

29. Vidal-Pineiro D, Sorensen O, Stromstad M, Amlien IK, Baare WFC, Bartres-Faz D, et al. Vulnerability to memory decline in aging. A mega-analysis of structural brain change. [Internet]. Neuroscience; 2025 [cited 2025 Apr 17]. Available from: http://biorxiv.org/lookup/doi/10.1101/2025.03.27.642988

30. Rehák Bučková B, Fraza C, Rehák R, Kolenič M, Beckmann C, Španiel F, et al. Using normative models pre-trained on cross-sectional data to evaluate intra-individual longitudinal changes in neuroimaging data [Internet]. 2025 [cited 2025 May 16]. Available from: https://elifesciences.org/reviewed-preprints/95823v3

31. Gaiser C, Berthet P, Kia SM, Frens MA, Beckmann CF, Muetzel RL, et al. Estimating cortical thickness trajectories in children across different scanners using transfer learning from normative models. Hum Brain Mapp. 2024;45(2):e26565.

32. Janssen J, Gallego AG, Díaz-Caneja CM, Lois NG, Janssen N, González-Peñas J, et al. Heterogeneity of morphometric similarity networks in health and schizophrenia [Internet]. Neuroscience; 2024 [cited 2025 May 16]. Available from: http://biorxiv.org/lookup/doi/10.1101/2024.03.26.586768

33. Bayer JMM, Dinga R, Kia SM, Kottaram AR, Wolfers T, Lv J, et al. Accommodating site variation in neuroimaging data using normative and hierarchical Bayesian models. NeuroImage. 2022 Dec;264:119699.

34. Dehestani N, Whittle S, Vijayakumar N, Silk TJ. Developmental brain changes during puberty and associations with mental health problems. Dev Cogn Neurosci. 2023 Apr;60:101227.

35. Rogers JC, De Brito SA. Cortical and Subcortical Gray Matter Volume in Youths With Conduct Problems: A Meta-analysis. JAMA Psychiatry. 2016 Jan 1;73(1):64.

36. Solmi M, Radua J, Olivola M, Croce E, Soardo L, Salazar De Pablo G, et al. Age at onset of mental disorders worldwide: large-scale meta-analysis of 192 epidemiological studies. Mol Psychiatry. 2022 Jan;27(1):281–95.

37. Whittle S, Vijayakumar N, Simmons JG, Allen NB. Internalizing and Externalizing Symptoms Are Associated With Different Trajectories of Cortical Development During Late Childhood. J Am Acad Child Adolesc Psychiatry. 2020 Jan;59(1):177–85.

38. Dehestani N, Vijayakumar N, Ball G, Mansour L S, Whittle S, Silk TJ. “Puberty age gap”: new method of assessing pubertal timing and its association with mental health problems. Mol Psychiatry. 2024 Feb;29(2):221–8.

39. Casey BJ, Cannonier T, Conley MI, Cohen AO, Barch DM, Heitzeg MM, et al. The Adolescent Brain Cognitive Development (ABCD) study: Imaging acquisition across 21 sites. Dev Cogn Neurosci. 2018 Aug;32:43–54.

40. Herting MM, Uban KA, Gonzalez MR, Baker FC, Kan EC, Thompson WK, et al. Correspondence Between Perceived Pubertal Development and Hormone Levels in 9-10 Year-Olds From the Adolescent Brain Cognitive Development Study. Front Endocrinol. 2021 Feb 18;11:549928.

41. Kraft D, Alnæs D, Kaufmann T. Domain adapted brain network fusion captures variance related to pubertal brain development and mental health. Nat Commun. 2023 Oct 23;14(1):6698.

42. Cheng TW, Magis-Weinberg L, Williamson VG, Ladouceur CD, Whittle SL, Herting MM, et al. A Researcher’s Guide to the Measurement and Modeling of Puberty in the ABCD Study® at Baseline. Front Endocrinol [Internet]. 2021 [cited 2025 Apr 14];12. Available from: https://www.readcube.com/articles/10.3389%2Ffendo.2021.608575

43. Petersen AC, Crockett L, Richards M, Boxer A. A self-report measure of pubertal status: Reliability, validity, and initial norms. J Youth Adolesc. 1988 Apr;17(2):117–33.

44. Fischl B. FreeSurfer. NeuroImage. 2012 Aug;62(2):774–81.

45. Glasser MF, Coalson TS, Robinson EC, Hacker CD, Harwell J, Yacoub E, et al. A multi-modal parcellation of human cerebral cortex. Nature. 2016 Aug;536(7615):171–8.

46. Laurent JS, Watts R, Adise S, Allgaier N, Chaarani B, Garavan H, et al. Associations Among Body Mass Index, Cortical Thickness, and Executive Function in Children. JAMA Pediatr. 2020 Feb 1;174(2):170.

47. Veit R, Kullmann S, Heni M, Machann J, Häring HU, Fritsche A, et al. Reduced cortical thickness associated with visceral fat and BMI. NeuroImage Clin. 2014;6:307–11.

48. Westwater ML, Vilar-López R, Ziauddeen H, Verdejo-García A, Fletcher PC. Combined effects of age and BMI are related to altered cortical thickness in adolescence and adulthood. Dev Cogn Neurosci. 2019 Dec;40:100728.

49. Rutherford S, Kia SM, Wolfers T, Fraza C, Zabihi M, Dinga R, et al. The normative modeling framework for computational psychiatry. Nat Protoc. 2022 July;17(7):1711–34.

50. Fraza CJ, Dinga R, Beckmann CF, Marquand AF. Warped Bayesian linear regression for normative modelling of big data. NeuroImage. 2021 Dec;245:118715.

51. Jones MC, Pewsey A. Sinh-arcsinh distributions. Biometrika. 2009 Dec 1;96(4):761–80.

52. Dinga R, Fraza CJ, Bayer JMM, Kia SM, Beckmann CF, Marquand AF. Normative modeling of neuroimaging data using generalized additive models of location scale and shape [Internet]. 2021 [cited 2025 Feb 1]. Available from: http://biorxiv.org/lookup/doi/10.1101/2021.06.14.448106

53. Benjamini Y, Hochberg Y. Controlling the False Discovery Rate: A Practical and Powerful Approach to Multiple Testing. J R Stat Soc Ser B Stat Methodol. 1995 Jan 1;57(1):289–300.

54. Kaufmann T, Karolinska Schizophrenia Project (KaSP), Van Der Meer D, Doan NT, Schwarz E, Lund MJ, et al. Common brain disorders are associated with heritable patterns of apparent aging of the brain. Nat Neurosci. 2019 Oct;22(10):1617–23.

55. Marquand A, Rutherford S, Dinga R. Fairly evaluating the performance of normative models. Lancet Digit Health. 2024 Nov;6(11):e775.

56. Aris IM, Rifas-Shiman SL, Zhang X, Yang S, Switkowski K, Fleisch AF, et al. Association of BMI with Linear Growth and Pubertal Development. Obesity. 2019 Oct;27(10):1661–70.

57. Bramen JE, Hranilovich JA, Dahl RE, Chen J, Rosso C, Forbes EE, et al. Sex Matters during Adolescence: Testosterone-Related Cortical Thickness Maturation Differs between Boys and Girls. Zang YF, editor. PLoS ONE. 2012 Mar 29;7(3):e33850.

58. Wierenga LM, Doucet GE, Dima D, Agartz I, Aghajani M, Akudjedu TN, et al. Greater male than female variability in regional brain structure across the lifespan. Hum Brain Mapp. 2022;43(1):470–99.

59. Beck D, Ferschmann L, MacSweeney N, Norbom LB, Wiker T, Aksnes E, et al. Puberty differentially predicts brain maturation in male and female youth: A longitudinal ABCD Study. Dev Cogn Neurosci. 2023 June;61:101261.

60. Brouwer RM, Koenis MMG, Schnack HG, Van Baal GC, Van Soelen ILC, Boomsma DI, et al. Longitudinal Development of Hormone Levels and Grey Matter Density in 9 and 12-Year-Old Twins. Behav Genet. 2015 May;45(3):313– 23.

61. Bashyam VM, Erus G, Doshi J, Habes M, Nasrallah IM, Truelove-Hill M, et al. MRI signatures of brain age and disease over the lifespan based on a deep brain network and 14 468 individuals worldwide. Brain. 2020 July 1;143(7):2312–24.

62. Hahn T, Fisch L, Ernsting J, Winter NR, Leenings R, Sarink K, et al. From ‘loose fitting’ to high-performance, uncertainty-aware brain-age modelling. Brain. 2021 Apr 12;144(3):e31–e31.

63. Shaw P, Kabani NJ, Lerch JP, Eckstrand K, Lenroot R, Gogtay N, et al. Neurodevelopmental Trajectories of the Human Cerebral Cortex. J Neurosci. 2008 Apr 2;28(14):3586–94.

64. Tamnes CK, Herting MM, Goddings AL, Meuwese R, Blakemore SJ, Dahl RE, et al. Development of the Cerebral Cortex across Adolescence: A Multisample Study of Inter-Related Longitudinal Changes in Cortical Volume, Surface Area, and Thickness. J Neurosci. 2017 Mar 22;37(12):3402–12.

65. Herting MM, Gautam P, Spielberg JM, Dahl RE, Sowell ER. A Longitudinal Study: Changes in Cortical Thickness and Surface Area during Pubertal Maturation. Baud O, editor. PLOS ONE. 2015 Mar 20;10(3):e0119774.

66. Vijayakumar N, Allen NB, Youssef G, Dennison M, Yücel M, Simmons JG, et al. Brain development during adolescence: A mixed-longitudinal investigation of cortical thickness, surface area, and volume. Hum Brain Mapp. 2016 June;37(6):2027–38.

67. Crone EA, Dahl RE. Understanding adolescence as a period of social–affective engagement and goal flexibility. Nat Rev Neurosci. 2012 Sept;13(9):636–50.

68. Mills KL, Goddings AL, Clasen LS, Giedd JN, Blakemore SJ. The Developmental Mismatch in Structural Brain Maturation during Adolescence. Dev Neurosci. 2014;36(3–4):147–60.

