## Supplementary Material for "Predicting Developmental Norms from Baseline Cortical Thickness in Longitudinal Studies"

### Figures

### Tables

### 1. Excluding participants using Euler numbers

**Training set: participants with only baseline- and 2-year follow-up data.**

A. **Before**

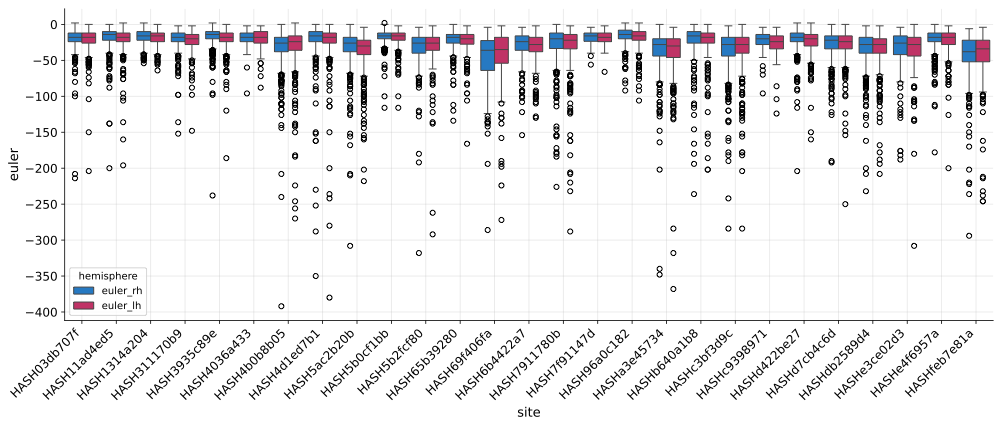

B. **After**

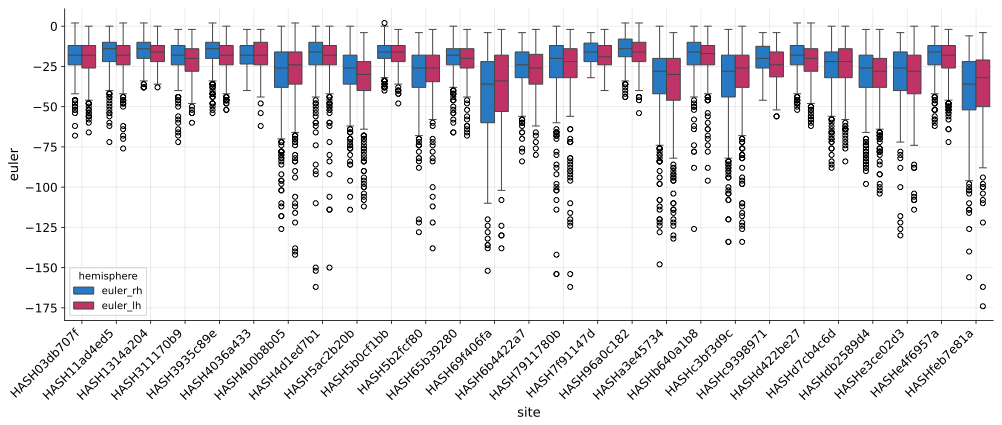

Figure S 1 Before and-after exclusion of badly reconstructed Freesurfer data based on Euler numbers: training data.

Mean and standard deviations were computed using Euler numbers for each site individually. Participants whose Euler numbers exceeded 6*SD from the mean were excluded.

**Test set: participants with baseline-, 2-year, and 4-year follow-up data.**

A. **Before**

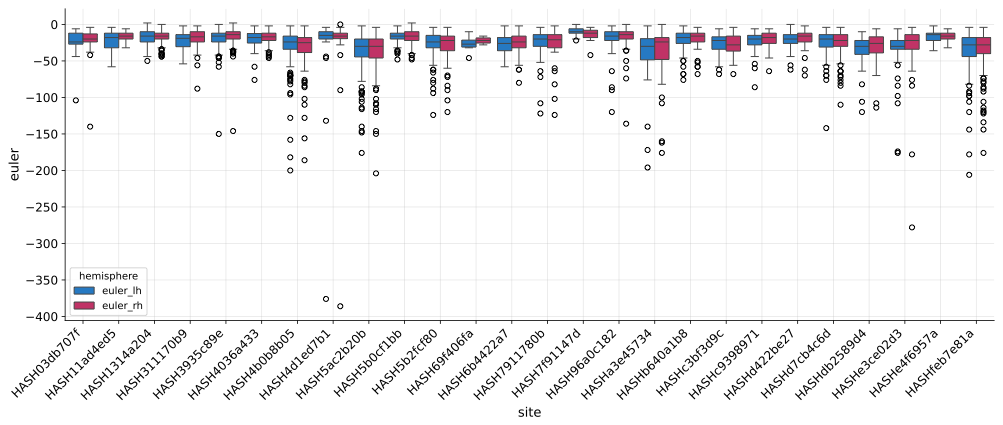

B. **After**

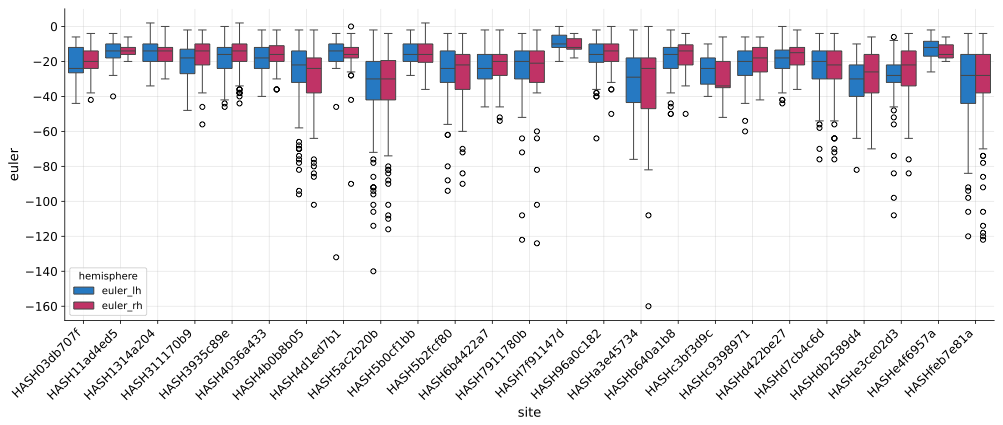

Figure S 2 Before and-after exclusion of badly reconstructed Freesurfer data based on Euler numbers: test data

Mean and standard deviations were computed using Euler numbers for each site individually. Participants whose Euler numbers exceeded 6*SD from the mean were excluded.

### 2. Sanity checks after preprocessing

A. **Age distributions**

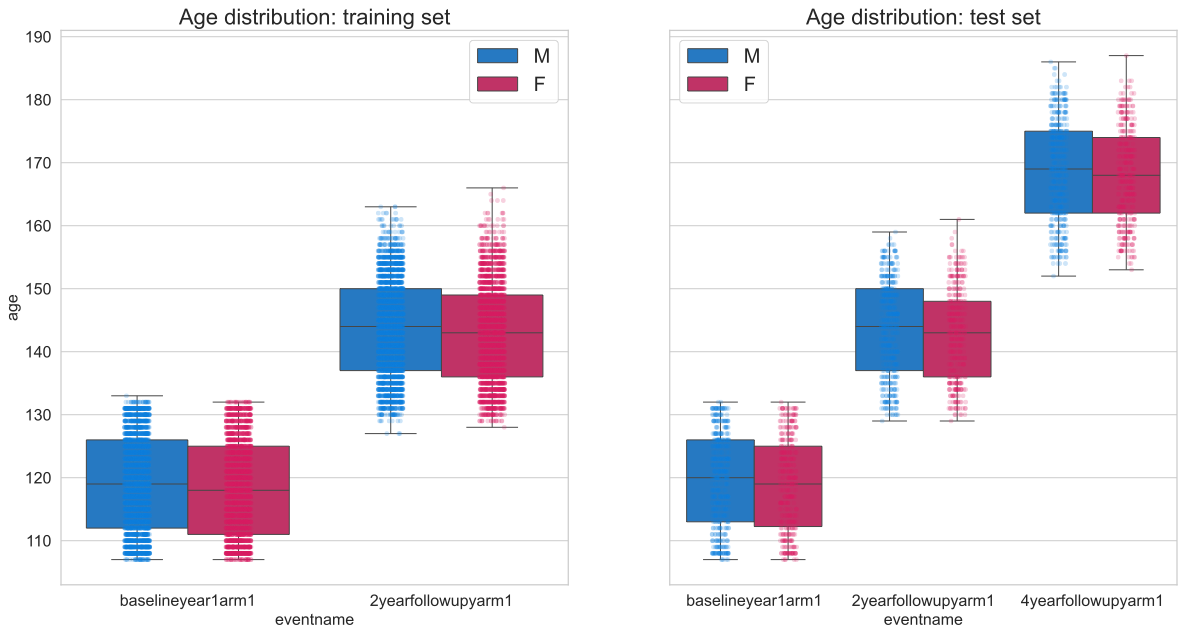

B. **BMI distributions**

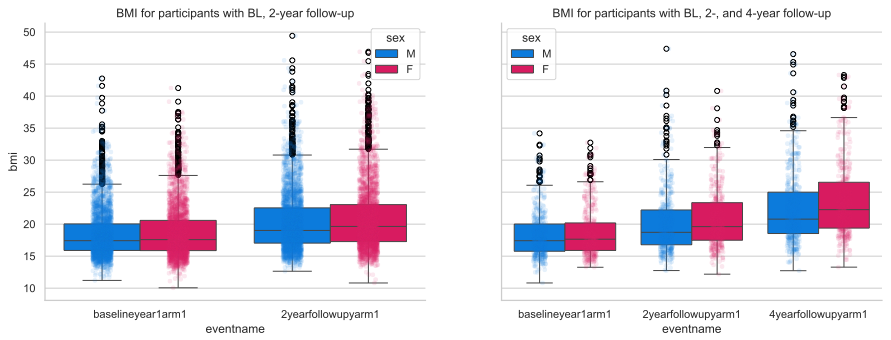

Figure S 3 Demographics for training and test set.

We found no discernable difference in age between males and females within timepoints (Panel A). Average body mass index increased slightly with age but was not statistically different between males and females.

We performed Welch’s t-tests to test for differences of baseline ages and BMI between the training and test dataset. These tests revealed no statistical differences for age (male: t(3522)=0.3876, p<0.7; female: t(3056)=-0.469, p<0.64) nor for BMI (male: t(3522)=-0.467, p<0.64; female: t(3056)=1.54, p<0.124).

**A. Average cortical thickness for the train and test data**

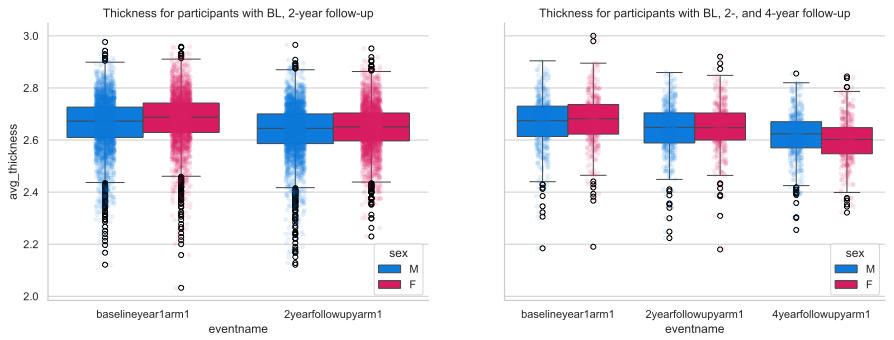

**B. Puberty scores as rated by caregivers for the train and test data**

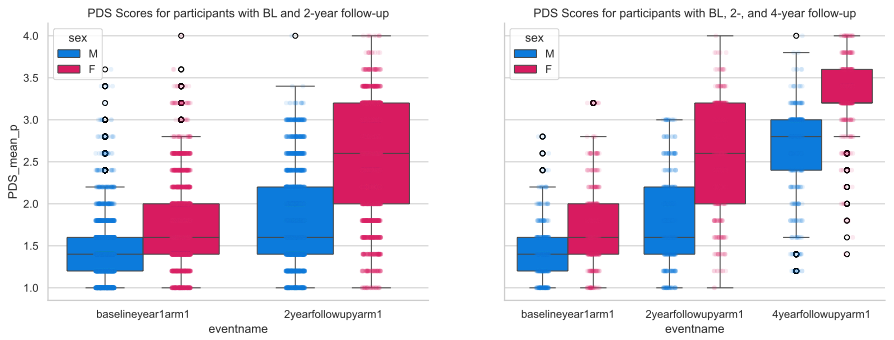

Figure S 4 Changes in cortical thickness and pubertal development across visits.

Panel A) As was expected, we saw a decline in cortical thickness with increasing age. This expected decline was also more pronounced in females than in males at 4-year follow-up. Panel B) illustrates the changes in the pubertal development scale (PDS) as rated by the youth’s caregivers over time. As expected, we observed a stronger increase for females than for males. Although it should be noted that the menarche item is binary (i.e., either 1 or 4), which may explain the dramatic increase over males in the 2-year follow-up data.

A Welch’s t-test revealed a statistical difference between the baseline PDS mean scores between the training and test set in both males (t(3522)=-9.47, p<4.7e^-21^) and females (t(3056)=1.568, p<6.49e^-14^).

**Distribution of males and females according to ethnicity**

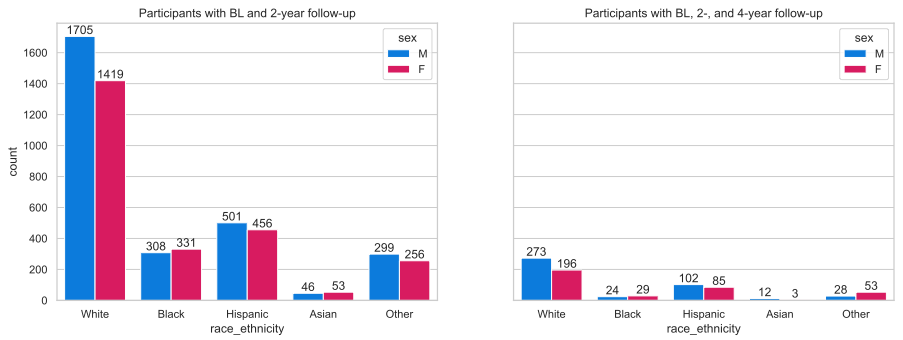

Figure S 5 Distribution of the ABCD ethnicities in the training (left) and test (right) datasets.

### 3. Supplementary information to results section: Longitudinal B-Norm models better predict cortical thickness at future time points than standard Age models

#### Mean standardized log loss (MSLL)

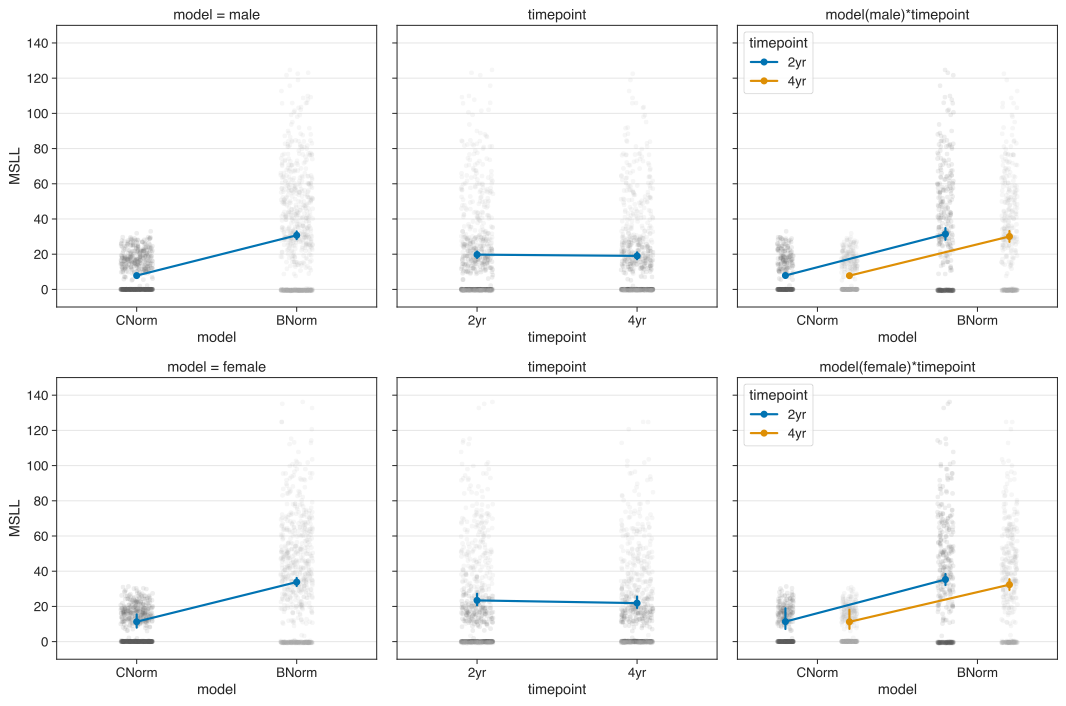

Figure S 6 Interaction plots for mean standardized log loss.

Note: We limited the y-axes to [-10, 150] as there were some extreme outliers that squished everything such that nothing was really visible.

Table S 1 Mean standardized log loss statistics

| ***model*** | ***sex*** | ***timepoint*** | ***mean*** | ***sem*** |
| --- | --- | --- | --- | --- |
| C-Norm | male | 2yr | 1,07E+10 | 6,33E+09 |
|  |  | 4yr | 1,04E+10 | 6,14E+09 |
|  | female | 2yr | 3,24E+10 | 1,59E+10 |
|  |  | 4yr | 3,07E+10 | 1,51E+10 |
| B-Norm | male | 2yr | 2,17E+09 | 1,57E+09 |
|  |  | 4yr | 2,05E+09 | 1,49E+09 |
|  | female | 2yr | 1,50E+09 | 9,36E+08 |
|  |  | 4yr | 1,39E+09 | 8,68E+08 |

Repeated measures ANOVA for male models:

|  | **sum_sq (III)** | **df** | **df2** | **F** | **PR(>F)** | **np2** |
| --- | --- | --- | --- | --- | --- | --- |
| model | 2.560779e+10 | 1.0 | 359 | 1.718 | 0.190697 | 0.004765 |
| timepoint | 2.265690e+07 | 1.0 | 359 | 5.457 | 0.020039 | 0.014973 |
| model×timepoint | 6.811265e+06 | 1.0 | 359 | 1.623 | 0.203473 | 0.004501 |

Pair-wise tests for male models:

|  | **model** | **A** | **B** | **T** | **dof** | **p-unc** | **hedges** |
| --- | --- | --- | --- | --- | --- | --- | --- |
| model | - | C-Norm | B-Norm | 1.310995 | 359 | 0.190697 | 0.097760 |
| timepoint | - | 2yr | 4yr | 2.336034 | 359 | 0.020039 | 0.004118 |
| model×timepoint | C-Norm | 2yr | 4yr | 1.947903 | 359 | 0.052206 | 0.003277 |
| model×timepoint | B-Norm | 2yr | 4yr | 1.393163 | 359 | 0.164433 | 0.003902 |

Repeated measures ANOVA for female models:

|  | **sum_sq (III)** | **df** | **df2** | **F** | **PR(>F)** | **np2** |
| --- | --- | --- | --- | --- | --- | --- |
| model | 3.263640e+11 | 1.0 | 359 | 3.746042 | 0.053718 | 0.010327 |
| timepoint | 2.687321e+08 | 1.0 | 359 | 4.658619 | 0.031560 | 0.012810 |
| model×timepoint | 2.031492e+08 | 1.0 | 359 | 3.510597 | 0.061790 | 0.009684 |

Pair-wise tests for female models:

|  | **model** | **A** | **B** | **T** | **dof** | **p-unc** | **hedges** |
| --- | --- | --- | --- | --- | --- | --- | --- |
| model | - | C-Norm | B-Norm | 1.935469 | 359 | 0.053718 | 0.144186 |
| timepoint | - | 2yr | 4yr | 2.158383 | 359 | 0.031560 | 0.005852 |
| model×timepoint | C-Norm | 2yr | 4yr | 2.023247 | 359 | 0.043788 | 0.005477 |
| model×timepoint | B-Norm | 2yr | 4yr | 1.654231 | 359 | 0.098955 | 0.006576 |

#### Explained Variance

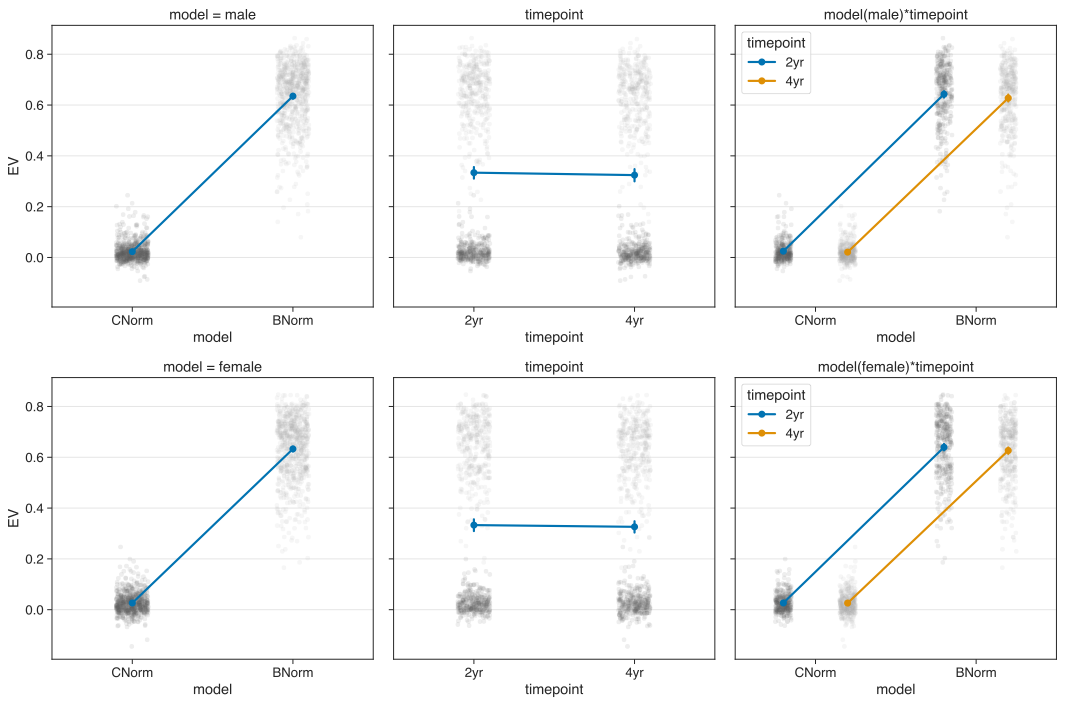

Figure S 7 Interaction plots for explained variance.

Table S 2 Explained variance statistics

| ***model*** | ***sex*** | ***timepoint*** | ***mean*** | ***sem*** |
| --- | --- | --- | --- | --- |
| C-Norm | male | 2yr | 0.024850 | 0.002010 |
|  |  | 4yr | 0.021023 | 0.002010 |
|  | female | 2yr | 0.026646 | 0.001956 |
|  |  | 4yr | 0.025996 | 0.002306 |
| B-Norm | male | 2yr | 0.642806 | 0.006439 |
|  |  | 4yr | 0.627440 | 0.006757 |
|  | female | 2yr | 0.639412 | 0.006537 |
|  |  | 4yr | 0.626474 | 0.006523 |

Repeated measures ANOVA for male models:

|  | **sum_sq (III)** | **df** | **df2** | **F** | **PR(>F)** | **np2** |
| --- | --- | --- | --- | --- | --- | --- |
| model | 134.918144 | 1.0 | 359 | 7855.030079 | 4.076e-246 | 0.956294 |
| timepoint | 0.033154 | 1.0 | 359 | 51.245127 | 4.633e-12 | 0.124913 |
| model×timepoint | 0.011983 | 1.0 | 359 | 25.864015 | 5.913e-07 | 0.067203 |

Pair-wise tests for male models:

|  | **model** | **A** | **B** | **T** | **dof** | **p-unc** | **hedges** |
| --- | --- | --- | --- | --- | --- | --- | --- |
| model | - | C-Norm | B-Norm | -88.628608 | 359 | 4.076e-246 | -6.71640 |
| timepoint | - | 2yr | 4yr | 7.158570 | 359 | 4.633e-12 | 0.148663 |
| model×timepoint | C-Norm | 2yr | 4yr | 3.267979 | 359 | 1.187e-03 | 0.100236 |
| model×timepoint | B-Norm | 2yr | 4yr | 7.015936 | 359 | 1.143e-11 | 0.122578 |

Repeated measures ANOVA for female models:

|  | **sum_sq (III)** | **df** | **df2** | **F** | **PR(>F)** | **np2** |
| --- | --- | --- | --- | --- | --- | --- |
| model | 132.47644 | 1.0 | 359 | 8803.735267 | 1.226e-254 | 0.960820 |
| timepoint | 0.016615 | 1.0 | 359 | 26.170219 | 5.100e-07 | 0.067945 |
| model×timepoint | 0.013590 | 1.0 | 359 | 26.040458 | 5.430e-07 | 0.067630 |

Pair-wise tests for female models:

|  | **model** | **A** | **B** | **T** | **dof** | **p-unc** | **hedges** |
| --- | --- | --- | --- | --- | --- | --- | --- |
| model | - | C-Norm | B-Norm | -93.828222 | 359 | 1.226e-254 | -6.68806 |
| timepoint | - | 2yr | 4yr | 5.115684 | 359 | 5.100e-07 | 0.099989 |
| model×timepoint | C-Norm | 2yr | 4yr | 0.612661 | 359 | 5.404e-01 | 0.015991 |
| model×timepoint | B-Norm | 2yr | 4yr | 5.618100 | 359 | 3.876e-08 | 0.104307 |

ROI-wise differences males:

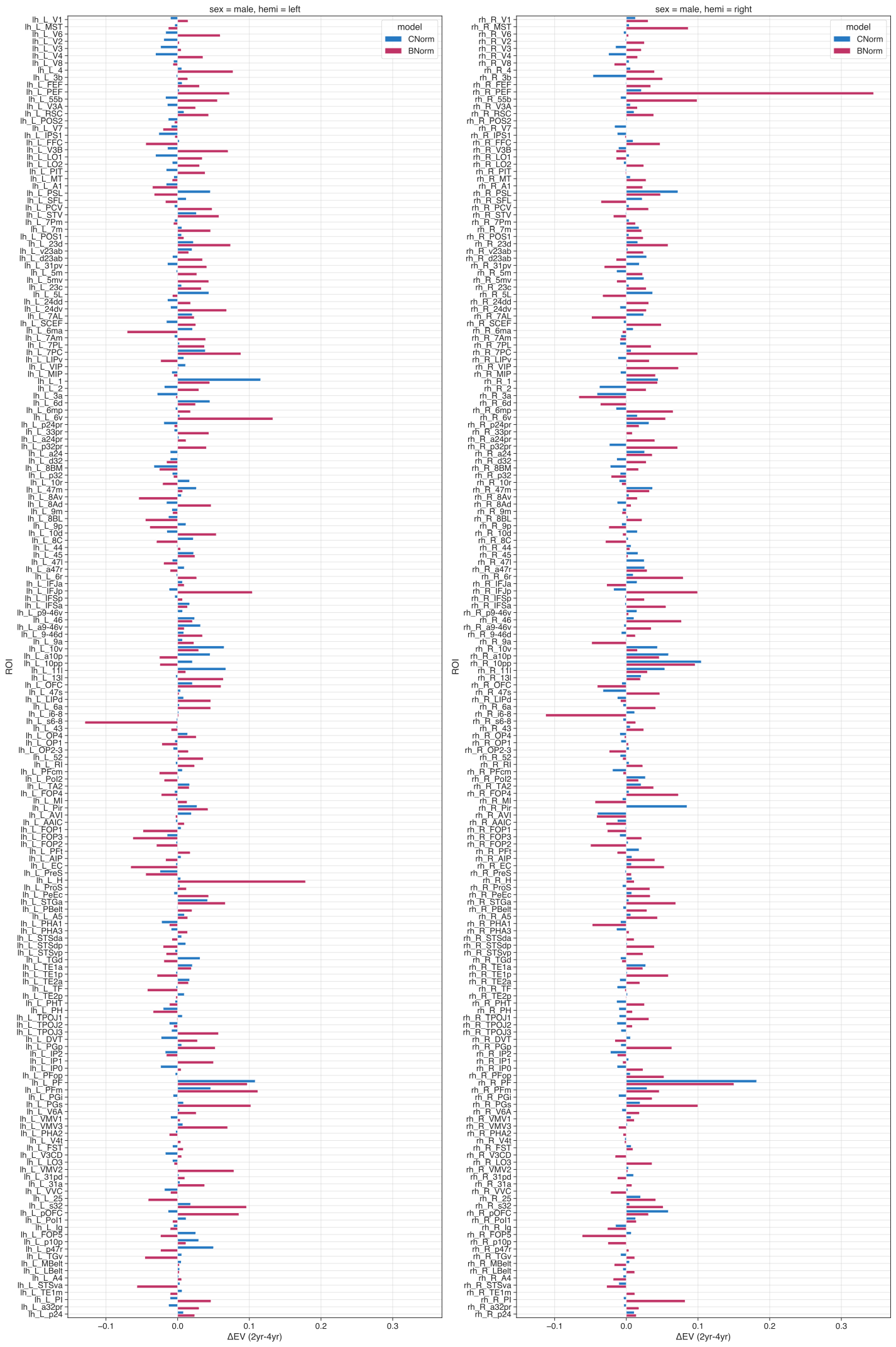

Figure S 8 ROI-wise difference (2year minus 4year) of explained variance for males.

ROI-wise differences females:

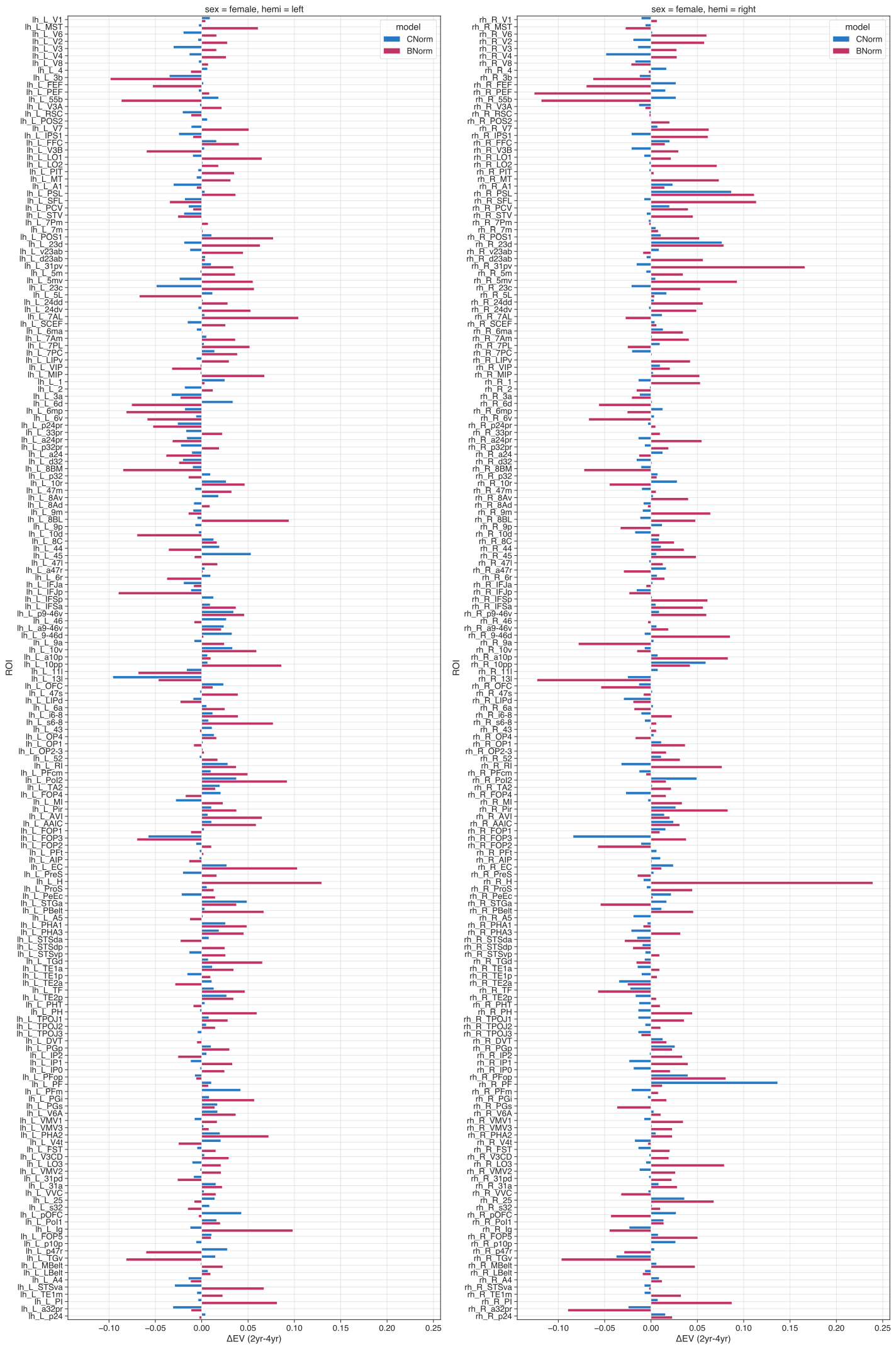

Figure S 9 ROI-wise difference (2year minus 4year) explained variance for females.

#### Standardized mean squared error (SMSE)

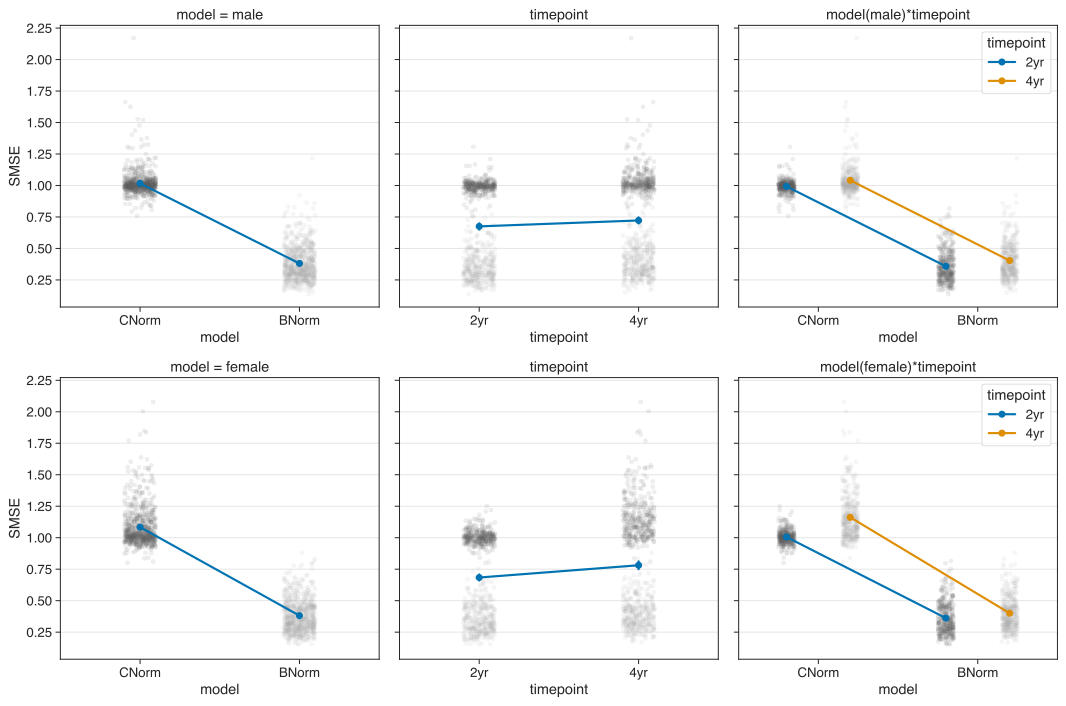

Figure S 10 Interaction plots for standardized mean squared error.

Table S 3 standardized mean squared error statistics

| ***model*** | ***sex*** | ***timepoint*** | ***mean*** | ***sem*** |
| --- | --- | --- | --- | --- |
| C-Norm | male | 2yr | 0.992541 | 0.002693 |
|  |  | 4yr | 1.041071 | 0.006351 |
|  | female | 2yr | 1.005893 | 0.002953 |
|  |  | 4yr | 1.162099 | 0.009572 |
| B-Norm | male | 2yr | 0.358037 | 0.006456 |
|  |  | 4yr | 0.403665 | 0.007461 |
|  | female | 2yr | 0.361767 | 0.006561 |
|  |  | 4yr | 0.400521 | 0.006735 |

Repeated measures ANOVA for male models:

|  | **sum_sq (III)** | **df** | **df2** | **F** | **PR(>F)** | **np2** |
| --- | --- | --- | --- | --- | --- | --- |
| model | 145.597865 | 1.0 | 359 | 7593.869315 | 1.347e-243 | 0.954859 |
| timepoint | 0.797900 | 1.0 | 359 | 261.395531 | 1.464e-44 | 0.421337 |
| model×timepoint | 0.000758 | 1.0 | 359 | 0.304499 | 5.814e-01 | 0.000847 |

Pair-wise tests for male models:

|  | **model** | **A** | **B** | **T** | **dof** | **p-unc** | **hedges** |
| --- | --- | --- | --- | --- | --- | --- | --- |
| model | - | C-Norm | B-Norm | 87.142810 | 359 | 1.347e-243 | 5.883945 |
| timepoint | - | 2yr | 4yr | -16.167731 | 359 | 1.464e-44 | -0.53849 |
| model×timepoint | C-Norm | 2yr | 4yr | -10.733332 | 359 | 1.703e-23 | -0.52384 |
| model×timepoint | B-Norm | 2yr | 4yr | -14.187876 | 359 | 1.389e-36 | -0.34434 |

Repeated measures ANOVA for female models:

|  | **sum_sq (III)** | **df** | **df2** | **F** | **PR(>F)** | **np2** |
| --- | --- | --- | --- | --- | --- | --- |
| model | 177.840354 | 1.0 | 359 | 7577.073462 | 1.969e-243 | 0.954764 |
| timepoint | 3.420820 | 1.0 | 359 | 602.006111 | 9.226e-79 | 0.626433 |
| model×timepoint | 1.241548 | 1.0 | 359 | 220.320024 | 3.370e-39 | 0.380308 |

Pair-wise tests for female models:

|  | **model** | **A** | **B** | **T** | **dof** | **p-unc** | **hedges** |
| --- | --- | --- | --- | --- | --- | --- | --- |
| model | - | C-Norm | B-Norm | 87.046387 | 359 | 1.969e-243 | 5.900791 |
| timepoint | - | 2yr | 4yr | -24.535813 | 359 | 9.226e-79 | -0.98817 |
| model×timepoint | C-Norm | 2yr | 4yr | -21.281107 | 359 | 1.364e-65 | -1.16110 |
| model×timepoint | B-Norm | 2yr | 4yr | -12.919172 | 359 | 1.280e-31 | -0.30688 |

#### Root mean squared error (RMSE)

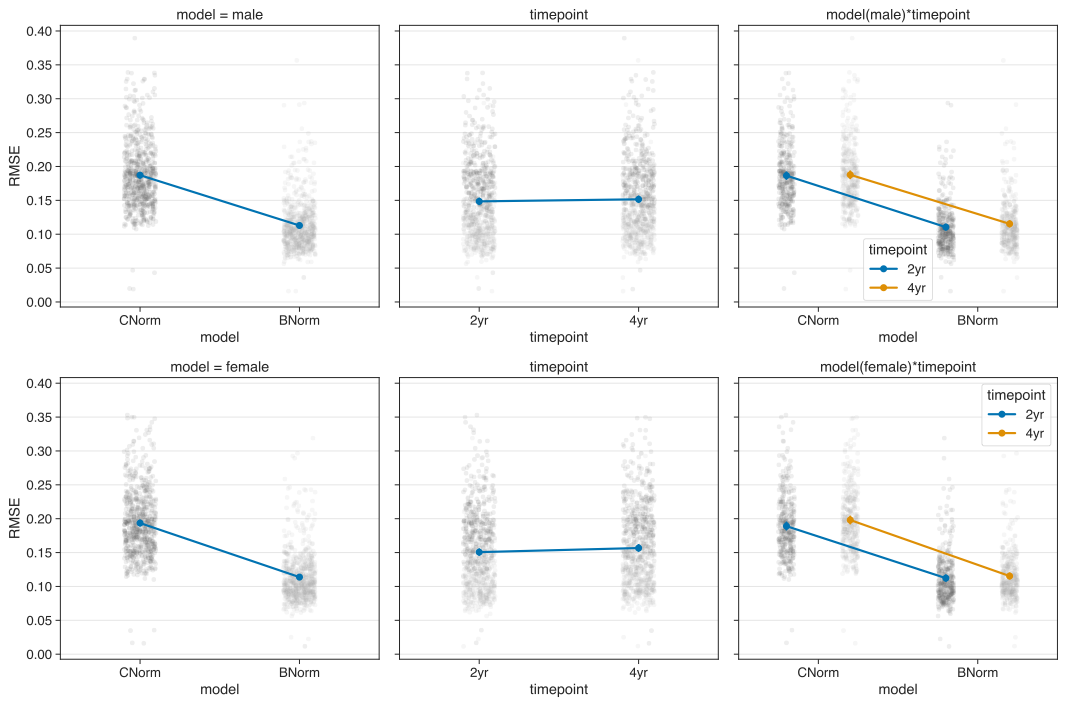

Figure S 11 Interaction plots for root mean squared error.

Table S 4 Root-mean squared error statistics

| ***model*** | ***sex*** | ***timepoint*** | ***mean*** | ***sem*** |
| --- | --- | --- | --- | --- |
| C-Norm | male | 2yr | 0.186527 | 0.002473 |
|  |  | 4yr | 0.197752 | 0.002589 |
|  | female | 2yr | 0.189054 | 0.002490 |
|  |  | 4yr | 0.198061 | 0.002544 |
| B-Norm | male | 2yr | 0.110464 | 0.001890 |
|  |  | 4yr | 0.115337 | 0.002007 |
|  | female | 2yr | 0.112266 | 0.002020 |
|  |  | 4yr | 0.115331 | 0.001892 |

Repeated measures ANOVA for male models:

|  | **sum_sq (III)** | **df** | **df2** | **F** | **PR(>F)** | **np2** |
| --- | --- | --- | --- | --- | --- | --- |
| model | 1.984116 | 1.0 | 359 | 2597.950827 | 1.883e-166 | 0.878591 |
| timepoint | 0.003347 | 1.0 | 359 | 61.269798 | 5.644e-14 | 0.145787 |
| model×timepoint | 0.001198 | 1.0 | 359 | 35.469328 | 6.177e-09 | 0.089917 |

Pair-wise tests for male models:

|  | **model** | **A** | **B** | **T** | **dof** | **p-unc** | **hedges** |
| --- | --- | --- | --- | --- | --- | --- | --- |
| model | - | C-Norm | B-Norm | 50.970097 | 359 | 1.883e-166 | 1.740543 |
| timepoint | - | 2yr | 4yr | -7.827503 | 359 | 5.644e-14 | -0.07525 |
| model×timepoint | C-Norm | 2yr | 4yr | -2.565330 | 359 | 1.071e-02 | -0.02547 |
| model×timepoint | B-Norm | 2yr | 4yr | -9.502101 | 359 | 3.011e-19 | -0.13162 |

Repeated measures ANOVA for female models:

|  | **sum_sq (III)** | **df** | **df2** | **F** | **PR(>F)** | **np2** |
| --- | --- | --- | --- | --- | --- | --- |
| model | 2.290157 | 1.0 | 359 | 3154.581077 | 6.678e-180 | 0.897825 |
| timepoint | 0.013117 | 1.0 | 359 | 169.514898 | 5.238e-32 | 0.320738 |
| model×timepoint | 0.003177 | 1.0 | 359 | 55.945537 | 5.778e-13 | 0.134826 |

Pair-wise tests for female models:

|  | **model** | **A** | **B** | **T** | **dof** | **p-unc** | **hedges** |
| --- | --- | --- | --- | --- | --- | --- | --- |
| model | - | C-Norm | B-Norm | 56.165657 | 359 | 6.678e-180 | 1.880078 |
| timepoint | - | 2yr | 4yr | -13.019789 | 359 | 5.238e-32 | -0.14917 |
| model×timepoint | C-Norm | 2yr | 4yr | -12.946247 | 359 | 1.006e-31 | -0.18840 |
| model×timepoint | B-Norm | 2yr | 4yr | -5.996011 | 359 | 4.931e-09 | -0.08248 |

#### Pearson’s correlation (real vs. predicted)

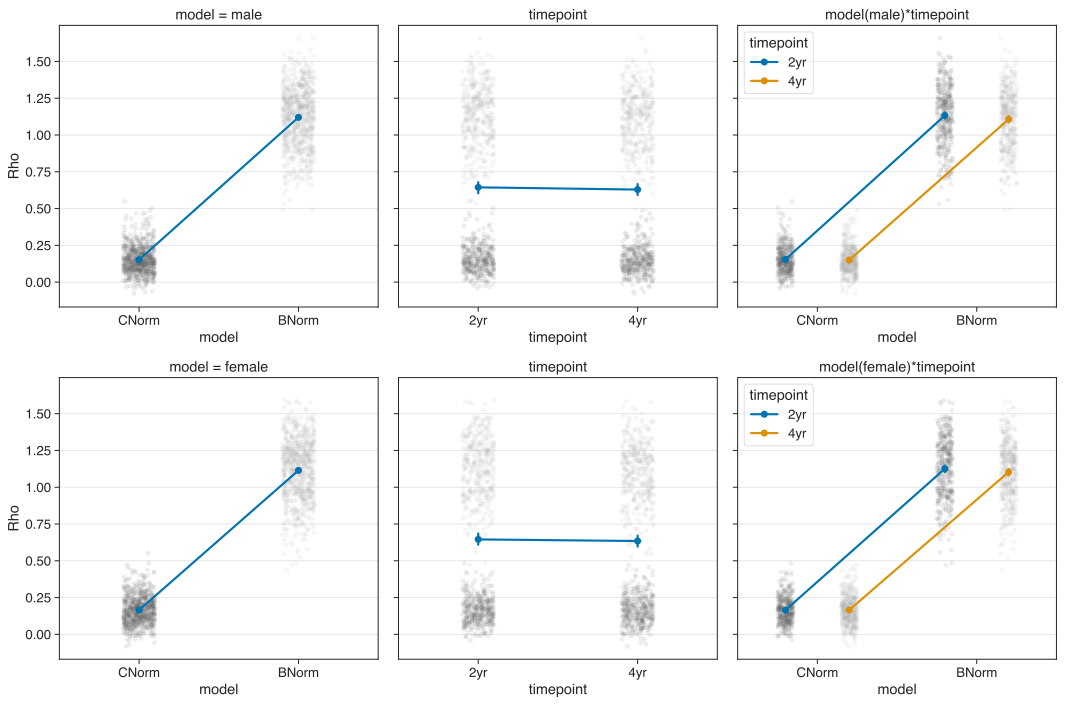

Figure S 12 Interaction plots for Pearson’s correlation.

Rho values were Fisher-z transformed before computing averages and before submitting to the rm-ANOVA.

Table S 5 Pearson’s rho statistics

| ***model*** | ***sex*** | ***timepoint*** | ***mean*** | ***sem*** |
| --- | --- | --- | --- | --- |
| C-Norm | male | 2yr | 0.151992 | 0.004744 |
|  |  | 4yr | 0.147164 | 0.004723 |
|  | female | 2yr | 0.162123 | 0.004587 |
|  |  | 4yr | 0.162319 | 0.005068 |
| B-Norm | male | 2yr | 0.800192 | 0.004052 |
|  |  | 4yr | 0.791132 | 0.004186 |
|  | female | 2yr | 0.796746 | 0.004320 |
|  |  | 4yr | 0.789628 | 0.004265 |

Repeated measures ANOVA for male models:

|  | **sum_sq (III)** | **df** | **df2** | **F** | **PR(>F)** | **np2** |
| --- | --- | --- | --- | --- | --- | --- |
| model | 149.712098 | 1.0 | 359 | 9634.944936 | 6.799e-261 | 0.964175 |
| timepoint | 0.017338 | 1.0 | 359 | 27.024021 | 3.385e-07 | 0.070188 |
| model×timepoint | 0.001617 | 1.0 | 359 | 3.275425 | 7.116e-02 | 0.009066 |

Pair-wise tests for male models:

|  | **model** | **A** | **B** | **T** | **dof** | **p-unc** | **hedges** |
| --- | --- | --- | --- | --- | --- | --- | --- |
| model | - | C-Norm | B-Norm | -98.157755 | 359 | 6.799e-261 | -7.82574 |
| timepoint | - | 2yr | 4yr | 5.198463 | 359 | 3.385e-07 | 0.125273 |
| model×timepoint | C-Norm | 2yr | 4yr | 2.255272 | 359 | 2.741e-02 | 0.053695 |
| model×timepoint | B-Norm | 2yr | 4yr | 6.872249 | 359 | 2.816e-11 | 0.116100 |

Repeated measures ANOVA for female models:

|  | **sum_sq (III)** | **df** | **df2** | **F** | **PR(>F)** | **np2** |
| --- | --- | --- | --- | --- | --- | --- |
| model | 143.322695 | 1.0 | 359 | 10563.04810 | 2.484e-268 | 0.967131 |
| timepoint | 0.004312 | 1.0 | 359 | 7.085329 | 8.121e-03 | 0.019354 |
| model×timepoint | 0.004814 | 1.0 | 359 | 9.475281 | 2.242e-03 | 0.025715 |

Pair-wise tests for female models:

|  | **model** | **A** | **B** | **T** | **dof** | **p-unc** | **hedges** |
| --- | --- | --- | --- | --- | --- | --- | --- |
| model | - | C-Norm | B-Norm | -102.77669 | 359 | 2.484e-268 | -7.40653 |
| timepoint | - | 2yr | 4yr | 2.661828 | 359 | 8.121e-03 | 0.054651 |
| model×timepoint | C-Norm | 2yr | 4yr | -0.096233 | 359 | 9.233e-01 | -0.00213 |
| model×timepoint | B-Norm | 2yr | 4yr | 4.958425 | 359 | 1.098e-06 | 0.087302 |

#### Skewness

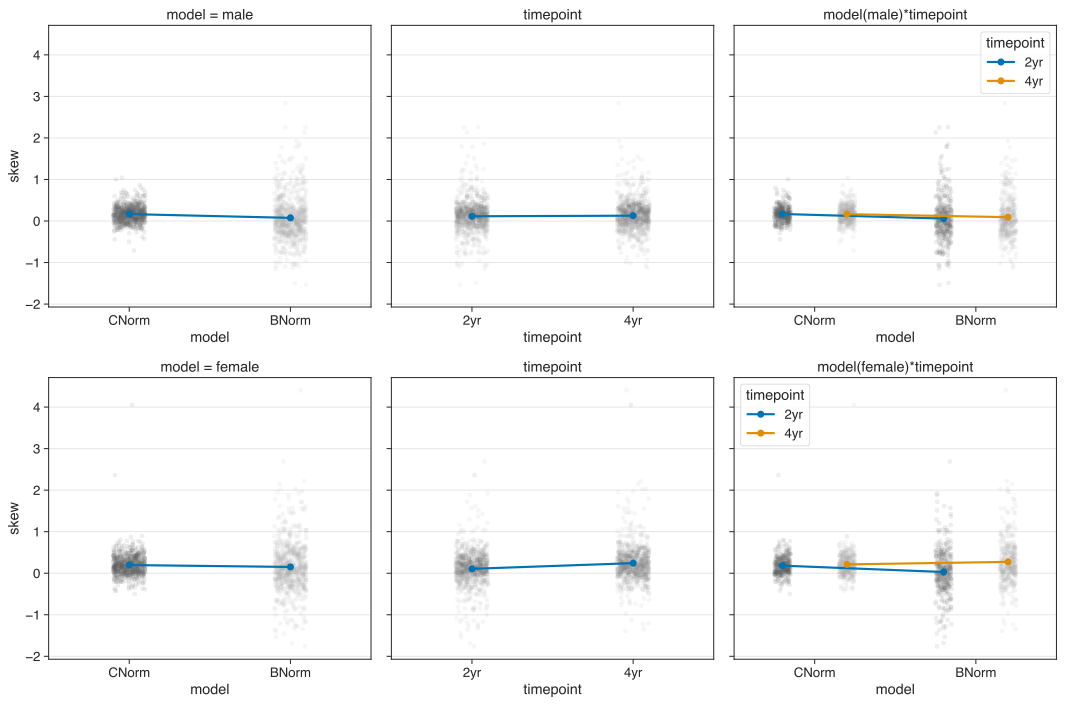

Figure S 13 Interaction plots for skewness.

We found more variance in skewness values for B-Norms (mean±SEM, 2-year: 0.0587±0.542, 4-year:0.092±0.497) than for C-Norms (2-year: 0.169±0.188, 4-year:0.165±0.209) in males. That is, our rm-ANOVA corroborated only the difference between the Age and longitudinal B-Norm models (sum of squares(SS)=2.997, F_1,359_=15.619, p<9.3e^-5^, $\eta^{2}$=0.041). For the female models, we found the lowest and highest average skewness for the 2-year follow-up data (0.029±0.573) and 4-year follow-up data (0.273±0.591) of the longitudinal *B-Norm* model, albeit again with larger variance than for the standard *Age* models (2-year=0.183±0.239; 4-year=0.213±0.302). As can be seen by the differential differences between the 2-year and 4-year follow-up in the longitudinal *B-Norm* models and the *Age* models, our rm-ANOVA revealed a statistically significant interaction effect of MODEL×TIMEPOINT (SS=4.134, F_1,359_=47.58, p<2.39e^-11^, $\eta^{2}$=0.117). Post-hoc tests are available in the supplements.

Table S 6 for skewness statistics

| ***model*** | ***sex*** | ***timepoint*** | ***mean*** | ***sem*** |
| --- | --- | --- | --- | --- |
| C-Norm | male | 2yr | 0.168878 | 0.009897 |
|  |  | 4yr | 0.164945 | 0.011034 |
|  | female | 2yr | 0.182953 | 0.012586 |
|  |  | 4yr | 0.212620 | 0.015939 |
| B-Norm | male | 2yr | 0.058715 | 0.028601 |
|  |  | 4yr | 0.092619 | 0.026188 |
|  | female | 2yr | 0.029373 | 0.030232 |
|  |  | 4yr | 0.273383 | 0.031180 |

Repeated measures ANOVA for male models:

|  | **sum_sq (III)** | **df** | **df2** | **F** | **PR(>F)** | **np2** |
| --- | --- | --- | --- | --- | --- | --- |
| model | 2.997198 | 1.0 | 359 | 15.618609 | 0.000093 | 0.041692 |
| timepoint | 0.080848 | 1.0 | 359 | 0.911024 | 0.340485 | 0.002531 |
| model×timepoint | 0.128851 | 1.0 | 359 | 2.072432 | 0.150854 | 0.005740 |

Pair-wise tests for male models:

|  | **model** | **A** | **B** | **T** | **dof** | **p-unc** | **hedges** |
| --- | --- | --- | --- | --- | --- | --- | --- |
| model | - | C-Norm | B-Norm | 3.952039 | 359 | 0.000093 | 0.266040 |
| timepoint | - | 2yr | 4yr | -0.954476 | 359 | 0.340485 | -0.04946 |
| model×timepoint | C-Norm | 2yr | 4yr | 0.570844 | 359 | 0.568462 | 0.019757 |
| model×timepoint | B-Norm | 2yr | 4yr | -1.205536 | 359 | 0.228790 | -0.06510 |

Repeated measures ANOVA for female models:

|  | **sum_sq (III)** | **df** | **df2** | **F** | **PR(>F)** | **np2** |
| --- | --- | --- | --- | --- | --- | --- |
| model | 0.775350 | 1.0 | 359 | 3.959325 | 4.737e-02 | 0.010908 |
| timepoint | 6.740953 | 1.0 | 359 | 43.002270 | 1.907e-10 | 0.106970 |
| model×timepoint | 4.134840 | 1.0 | 359 | 47.583383 | 2.389e-11 | 0.117032 |

Pair-wise tests for female models:

|  | **model** | **A** | **B** | **T** | **dof** | **p-unc** | **hedges** |
| --- | --- | --- | --- | --- | --- | --- | --- |
| model | - | C-Norm | B-Norm | 1.989805 | 359 | 4.737e-02 | 0.121346 |
| timepoint | - | 2yr | 4yr | -6.55761 | 359 | 1.907e-10 | -0.37040 |
| model×timepoint | C-Norm | 2yr | 4yr | -2.71135 | 359 | 7.022e-03 | -0.10876 |
| model×timepoint | B-Norm | 2yr | 4yr | -6.94649 | 359 | 1.767e-11 | -0.41834 |

ROI-wise differences males:

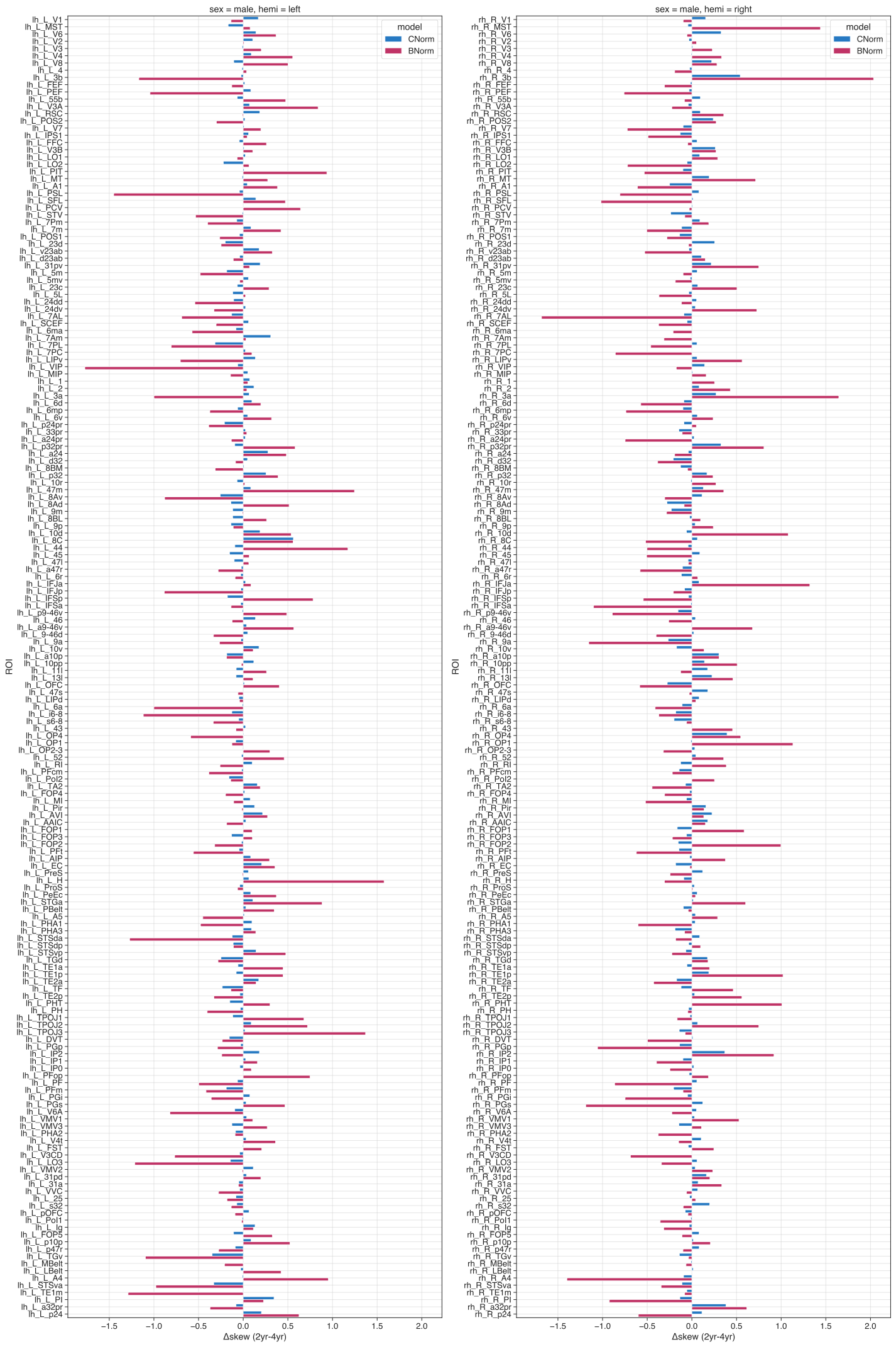

Figure S 14 ROI-wise difference (2year minus 4year) for skewness for males.

ROI-wise differences females:

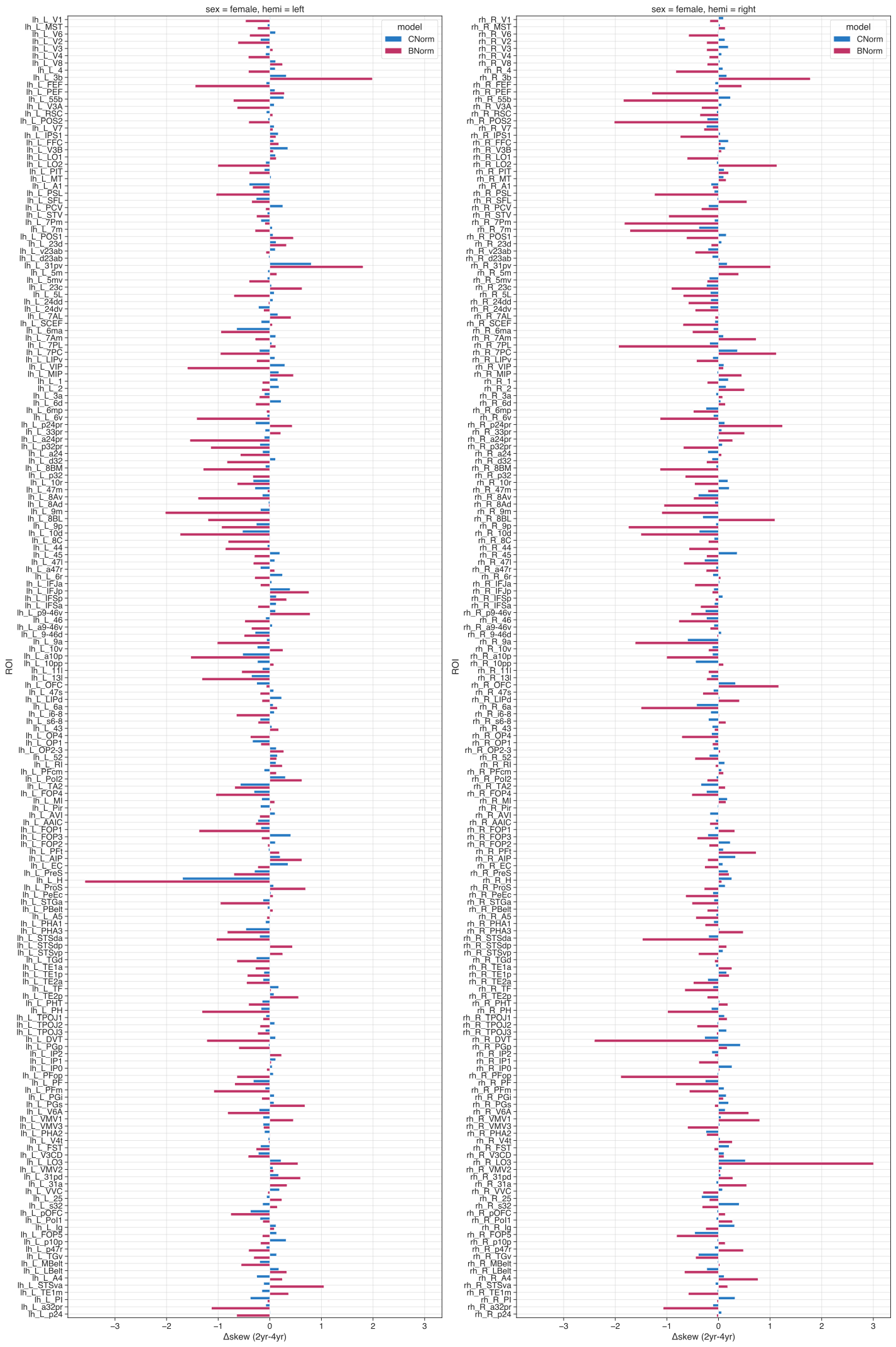

Figure S 15 ROI-wise difference (2year minus 4year) for skewness for females

#### Kurtosis

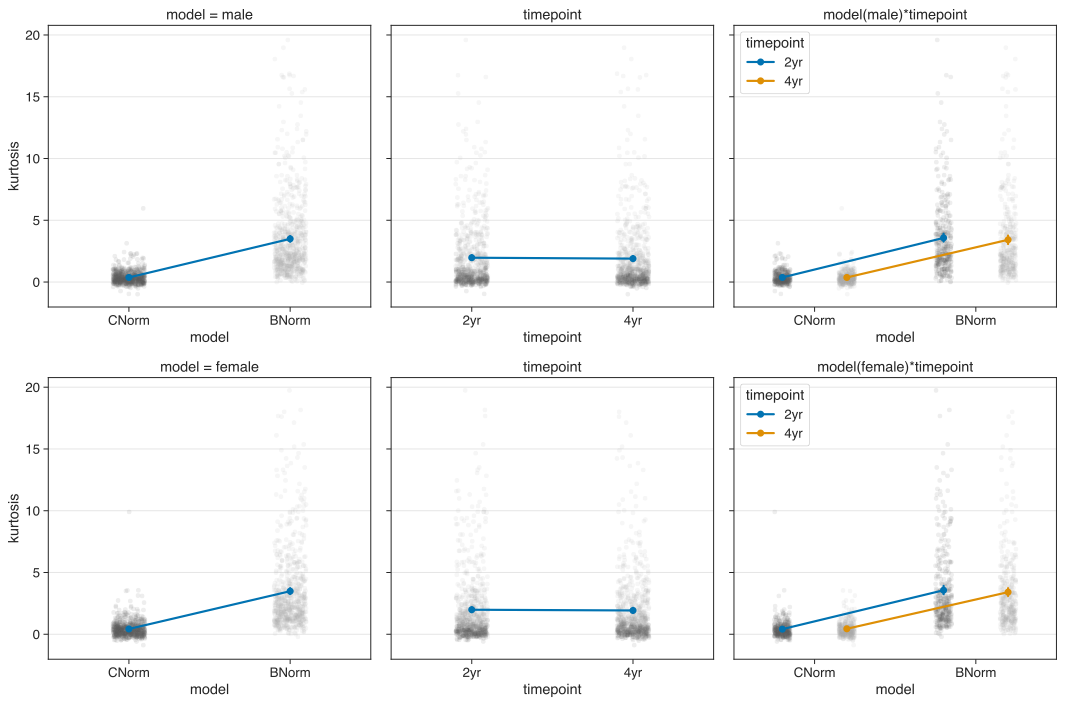

Figure S 16 Interactionplots for kurtosis.

As can be easily seen in the last row in panel A of Figure 1 of the main text, excess kurtosis was generally much larger in the longitudinal *B-Norm* models than in the *C-Norm* models, irrespective of sex and timepoint. We obtained lower kurtosis for the longitudinal *B-Norm* models in the 4-year follow-up data of the males compared to the 2-year follow-up data (2year-4year=0.2044).. This effect is somewhat reverse in the female Age models (2year-4year=-0.127). However, our rm-ANOVAs revealed no statistical effect of TIMEPOINT (males: SS=3.828, F_1,359_=2.241, p<0.135, $\eta^{2}$=0.0062; females: SS=0.298, F_1,359_=0.046, p<0.829, $\eta^{2}$=0.00013) or the MODEL×TIMEPOINT interaction (males: SS=3.69, F_1,359_=2.36, p<0.125, $\eta^{2}$=0.0065; females: SS=3.45, F_1,359_=1.462, p<0.227, $\eta^{2}$=0.00406).

Table S 7 for Kurtosis Statistics

| ***model*** | ***sex*** | ***timepoint*** | ***mean*** | ***sem*** |
| --- | --- | --- | --- | --- |
| C-Norm | male | 2yr | 0.371731 | 0.026005 |
|  |  | 4yr | 0.369859 | 0.028927 |
|  | female | 2yr | 0.414015 | 0.038817 |
|  |  | 4yr | 0.541394 | 0.102754 |
| B-Norm | male | 2yr | 3.624436 | 0.166718 |
|  |  | 4yr | 3.420059 | 0.175922 |
|  | female | 2yr | 3.610992 | 0.177697 |
|  |  | 4yr | 3.541184 | 0.211235 |

Repeated measures ANOVA for male models:

|  | **sum_sq (III)** | **df** | **df2** | **F** | **PR(>F)** | **np2** |
| --- | --- | --- | --- | --- | --- | --- |
| model | 3575.394208 | 1.0 | 359 | 403.233441 | 1.163e-60 | 0.529016 |
| timepoint | 3.828481 | 1.0 | 359 | 2.241535 | 1.352e-01 | 0.006205 |
| model×timepoint | 3.690724 | 1.0 | 359 | 2.364916 | 1.249e-01 | 0.006544 |

Pair-wise tests for male models:

|  | **model** | **A** | **B** | **T** | **dof** | **p-unc** | **hedges** |
| --- | --- | --- | --- | --- | --- | --- | --- |
| model | - | C-Norm | B-Norm | -20.080673 | 359 | 1.163e-60 | -1.46689 |
| timepoint | - | 2yr | 4yr | 1.497176 | 359 | 1.352e-01 | 0.061387 |
| model×timepoint | C-Norm | 2yr | 4yr | 0.080621 | 359 | 9.357e-01 | 0.003584 |
| model×timepoint | B-Norm | 2yr | 4yr | 1.539696 | 359 | 1.245e-01 | 0.062786 |

Repeated measures ANOVA for female models:

|  | **sum_sq (III)** | **df** | **df2** | **F** | **PR(>F)** | **np2** |
| --- | --- | --- | --- | --- | --- | --- |
| model | 3455.992461 | 1.0 | 359 | 419.236080 | 2.769e-62 | 0.538700 |
| timepoint | 0.298301 | 1.0 | 359 | 0.046851 | 8.287e-01 | 0.000130 |
| model×timepoint | 3.499407 | 1.0 | 359 | 1.462770 | 2.272e-01 | 0.004058 |

Pair-wise tests for female models:

|  | **model** | **A** | **B** | **T** | **dof** | **p-unc** | **hedges** |
| --- | --- | --- | --- | --- | --- | --- | --- |
| model | - | C-Norm | B-Norm | -20.475255 | 359 | 2.769e-62 | -1.29035 |
| timepoint | - | 2yr | 4yr | -0.216451 | 359 | 8.287e-01 | -0.01250 |
| model×timepoint | C-Norm | 2yr | 4yr | -1.645627 | 359 | 1.007e-01 | -0.08634 |
| model×timepoint | B-Norm | 2yr | 4yr | 0.337934 | 359 | 7.356e-01 | 0.018830 |

ROI-wise differences males:

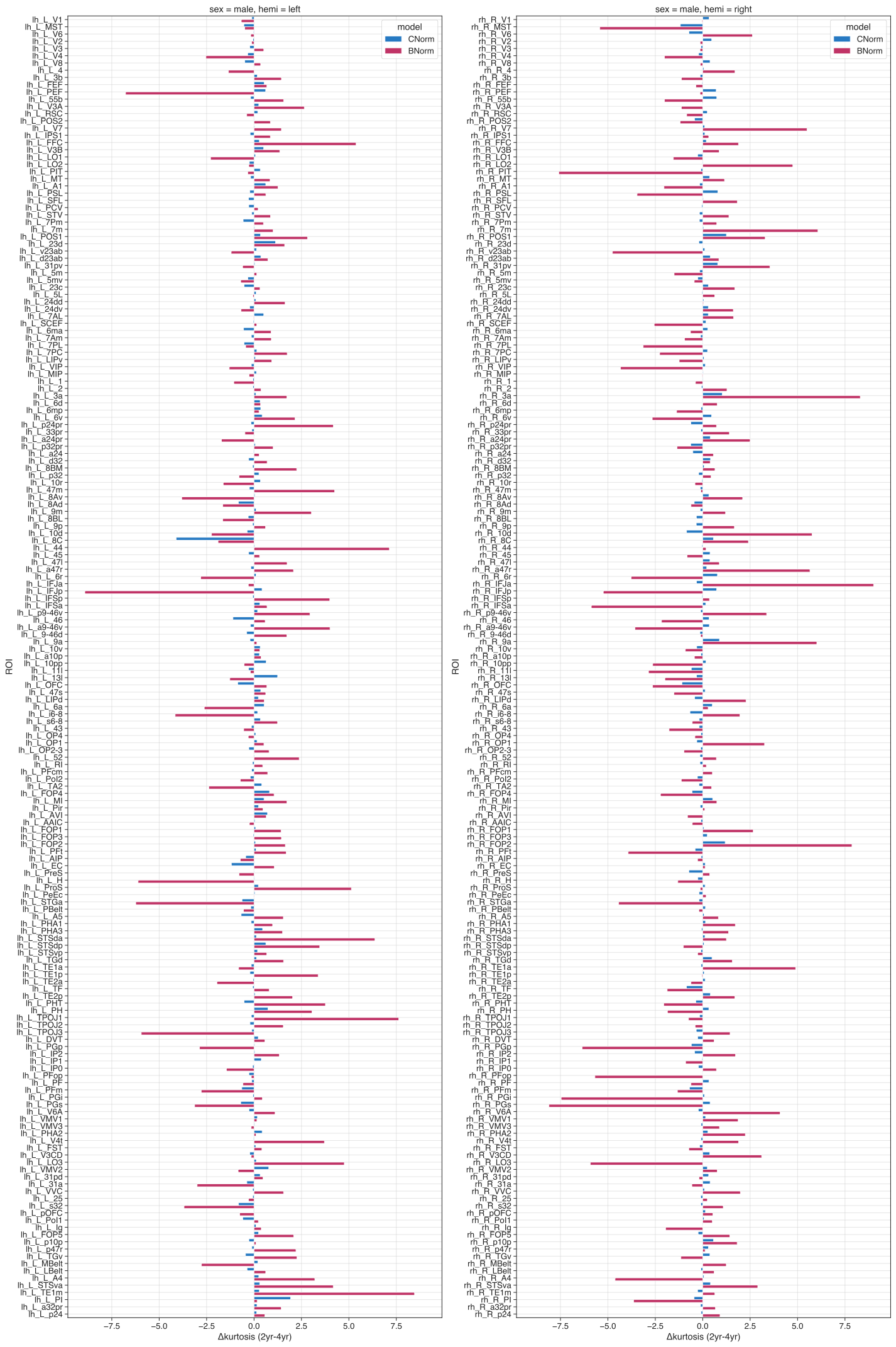

Figure S 17 ROI-wise difference (2year minus 4year) for kurtosis for males

ROI-wise differences females:

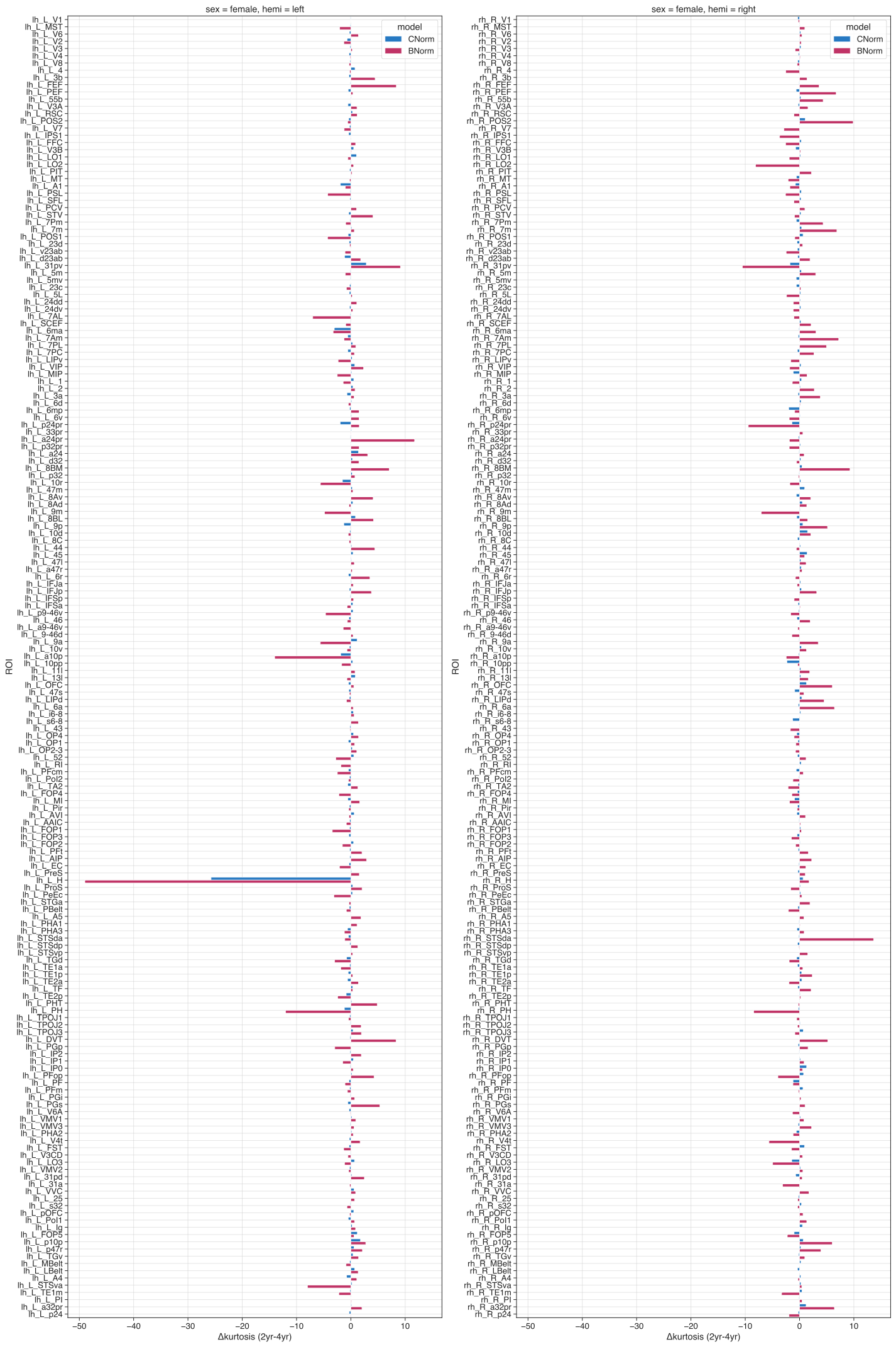

Figure S 18 ROI-wise difference (2year minus 4year) for kurtosis for females

#### Bayesian Information Criteria (BIC)

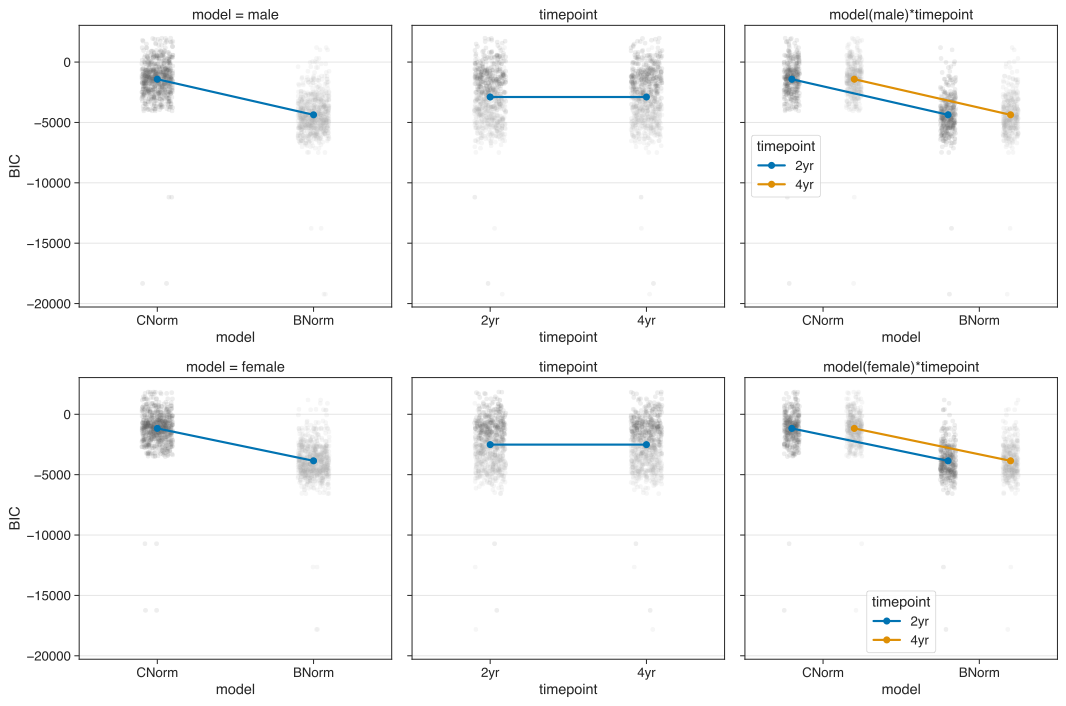

Figure S 19 Interaction plots for Bayesian Information Criteria (BIC).

Table S 8 for BIC Statistics

| ***model*** | ***sex*** | ***timepoint*** | ***mean*** | ***sem*** |
| --- | --- | --- | --- | --- |
| C-Norm | male | 2yr | -1418.224590 | 86.843136 |
|  |  | 4yr | -1418.224590 | 86.843136 |
|  | female | 2yr | -1168.022447 | 77.588938 |
|  |  | 4yr | -1168.022447 | 77.588938 |
| B-Norm | male | 2yr | -4368.158769 | 92.116311 |
|  |  | 4yr | -4368.158769 | 92.116311 |
|  | female | 2yr | -3853.867853 | 82.764235 |
|  |  | 4yr | -3853.867853 | 82.764235 |

Repeated measures ANOVA for male models:

|  | **sum_sq (III)** | **df** | **df2** | **F** | **PR(>F)** | **np2** |
| --- | --- | --- | --- | --- | --- | --- |
| model | 3.132760e+09 | 1.0 | 359 | 4423.606452 | 6.033e-204 | 0.924936 |
| timepoint | 0 | 1.0 | 359 | NaN | NaN | NaN |
| model×timepoint | 0 | 1.0 | 359 | NaN | NaN | NaN |

Pair-wise tests for male models:

|  | **model** | **A** | **B** | **T** | **dof** | **p-unc** | **hedges** |
| --- | --- | --- | --- | --- | --- | --- | --- |
| model | - | C-Norm | B-Norm | 66.510198 | 359 | 6.033e-204 | 1.734978 |
| timepoint | - | 2yr | 4yr | NaN | 359 | NaN | NaN |
| model×timepoint | C-Norm | 2yr | 4yr | NaN | 359 | NaN | NaN |
| model×timepoint | B-Norm | 2yr | 4yr | NaN | 359 | NaN | NaN |

Repeated measures ANOVA for female models:

|  | **sum_sq (III)** | **df** | **df2** | **F** | **PR(>F)** | **np2** |
| --- | --- | --- | --- | --- | --- | --- |
| model | 2.596956e+09 | 1.0 | 359 | 4435.001523 | 3.935e-204 | 0.925115 |
| timepoint | 0 | 1.0 | 359 | NaN | NaN | NaN |
| model×timepoint | 4.768372e-07 | 1.0 | 359 | -359.00 | 1 | -inf |

Pair-wise tests for female models:

|  | **model** | **A** | **B** | **T** | **dof** | **p-unc** | **hedges** |
| --- | --- | --- | --- | --- | --- | --- | --- |
| model | - | C-Norm | B-Norm | 66.595807 | 359 | 3.935e-204 | 1.762796 |
| timepoint | - | 2yr | 4yr | NaN | 359 | NaN | NaN |
| model×timepoint | C-Norm | 2yr | 4yr | NaN | 359 | NaN | NaN |
| model×timepoint | B-Norm | 2yr | 4yr | NaN | 359 | NaN | NaN |

#### Longitudinal normative plots for other baseline ages

| A Baseline age 114 months  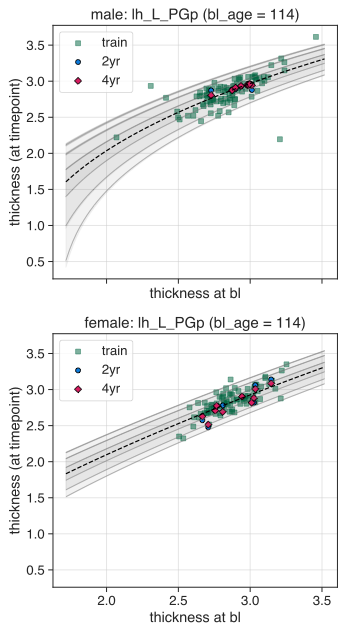 | B Baseline age 120 months  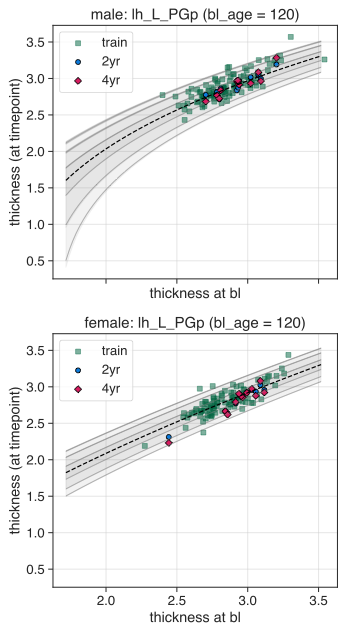 |
| --- | --- |
| Figure S 20 Normative plots of additional baseline ages.  For each baseline age the B-Norm models require to generate their own specific normative plots. | |

#### Surface maps for males and 4-year females

**Males:**

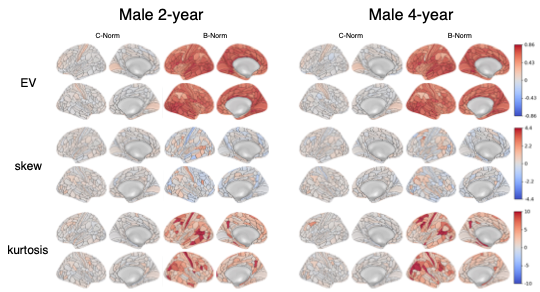

Figure S 21 ROI-wise performances for male C-Norms or B-Norms of 2-year and 4-year data.

The first two columns correspond to the lateral and medial views of the C-Norms, the last two to the B-Norms for the male data of the 2-year (left) and 4-year (right) timepoints. Warmer colors indicate more explained variance, positive skewness or kurtosis; colder colors indicate negative skewness or kurtosis.

**Females:**

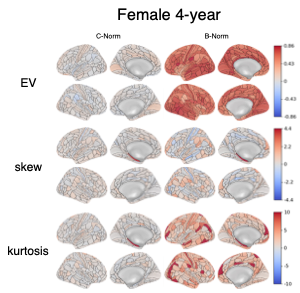

Figure S 22 ROI-wise performances for female C-Norms or B-Norms of 4-year data.

The first two columns correspond to the lateral and medial views of the C-Norms, the last two to the B-Norms for the female data of the 4-year (right) timepoint. Warmer colors indicate more explained variance, positive skewness or kurtosis; colder colors indicate negative skewness or kurtosis.

Table S 9 Kurtosis values greater 10 in the 2- or 4-year follow-up test data for both males and females and the cross-sectional normative models (C-Norms) or longitudinal normative models (B-Norms).

|  |  |  | TIMEPOINTS  (KURTOSIS) | |
| --- | --- | --- | --- | --- |
| ROI | ***sex*** | ***model*** | ***2yr*** | ***4yr*** |
| lh_L_31pv | female | B-Norm | 11,76290842 | 2,629781958 |
| lh_L_3b | female | B-Norm | 10,71668255 | 6,247540803 |
|  | male | B-Norm | 12,96071984 | 11,51556182 |
| lh_L_7AL | female | B-Norm | 3,8794104 | 10,90371176 |
| lh_L_7m | male | B-Norm | 16,5975464 | 15,59198511 |
| lh_L_8Av | male | B-Norm | 12,75218943 | 16,56447492 |
| lh_L_9a | female | B-Norm | 8,682803201 | 14,30709845 |
| lh_L_9m | female | B-Norm | 9,373430277 | 14,231347 |
| lh_L_DVT | female | B-Norm | 10,33011033 | 2,00732272 |
| lh_L_FEF | female | B-Norm | 11,31489956 | 2,940819297 |
| lh_L_FFC | male | B-Norm | 10,91524634 | 5,536031908 |
| lh_L_H | female | C-Norm | 9,901031485 | 35,64663669 |
|  |  | B-Norm | 1,603037042 | 50,58707693 |
|  | male | B-Norm | 4,75407098 | 10,86558246 |
| lh_L_IFJp | male | B-Norm | 1,148044584 | 10,06856346 |
| lh_L_PEF | male | B-Norm | 12,19898004 | 18,97121604 |
| lh_L_PGp | female | B-Norm | 14,65411217 | 17,62042069 |
| lh_L_PGs | female | B-Norm | 15,368533 | 10,03059332 |
| lh_L_PH | female | B-Norm | 1,598953809 | 13,62981184 |
| lh_L_PHA2 | male | B-Norm | 12,39759022 | 12,29477535 |
| lh_L_PHT | female | B-Norm | 10,00917361 | 5,149083075 |
| lh_L_PHT | male | B-Norm | 10,46510435 | 6,701237315 |
| lh_L_STGa | female | B-Norm | 13,50325382 | 13,848643 |
| lh_L_STGa | male | B-Norm | 5,76363991 | 11,99852562 |
| lh_L_STV | male | B-Norm | 11,3317584 | 10,4647438 |
| lh_L_TE1m | male | B-Norm | 11,49994023 | 3,038578396 |
| lh_L_TPOJ1 | male | B-Norm | 21,84947433 | 14,22164634 |
| lh_L_TPOJ2 | male | B-Norm | 19,59594667 | 18,05174349 |
| lh_L_V4t | male | B-Norm | 15,26645785 | 11,55654298 |
| lh_L_a10p | female | B-Norm | 1,152343017 | 15,18718302 |
| lh_L_a24pr | female | B-Norm | 13,31379949 | 1,574931428 |
| lh_L_a9-46v | female | B-Norm | 9,597011301 | 11,01125166 |
| lh_L_i6-8 | male | B-Norm | 9,296803167 | 13,45465258 |
| rh_R_10d | male | B-Norm | 10,24066773 | 4,475485798 |
| rh_R_31pv | female | B-Norm | 1,025028563 | 11,54680956 |
| rh_R_3a | male | B-Norm | 11,95140229 | 3,644869905 |
| rh_R_3b | female | B-Norm | 10,37805138 | 8,997568492 |
|  | male | B-Norm | 14,52870512 | 15,6496683 |
| rh_R_44 | female | B-Norm | 11,58053834 | 12,17725787 |
| rh_R_7Am | female | B-Norm | 20,5133387 | 13,30165131 |
| rh_R_7PL | female | B-Norm | 12,45543568 | 7,488820253 |
| rh_R_7m | female | B-Norm | 10,99658638 | 4,125820033 |
| rh_R_9a | female | B-Norm | 11,62209824 | 8,178718571 |
| rh_R_9m | female | B-Norm | 4,09637938 | 11,14340854 |
| rh_R_A4 | male | B-Norm | 5,586119388 | 10,20693123 |
| rh_R_DVT | female | B-Norm | 11,28552972 | 6,078333548 |
| rh_R_FOP2 | male | B-Norm | 10,05838288 | 2,190422599 |
| rh_R_IFJa | male | B-Norm | 16,74052749 | 7,729798258 |
| rh_R_LO2 | male | B-Norm | 10,78944139 | 6,03952023 |
| rh_R_LO3 | female | B-Norm | 13,05455982 | 18,00750931 |
|  | male | B-Norm | 4,249035663 | 10,17218253 |
| rh_R_MIP | female | B-Norm | 10,11372778 | 8,724604533 |
| rh_R_PFop | female | B-Norm | 6,515904078 | 10,50034521 |
| rh_R_PGi | male | B-Norm | 7,918783911 | 15,38134528 |
| rh_R_PGp | female | B-Norm | 17,66814705 | 16,1002063 |
|  | male | B-Norm | 10,51179197 | 16,86733645 |
| rh_R_PGs | female | B-Norm | 18,15842994 | 17,13469829 |
|  | male | B-Norm | 8,571392677 | 16,6776718 |
| rh_R_PH | female | B-Norm | 6,453859626 | 14,89600601 |
| rh_R_PIT | male | B-Norm | 9,164982121 | 16,75270585 |
| rh_R_POS2 | female | B-Norm | 12,82487944 | 2,946550829 |
| rh_R_PSL | male | B-Norm | 7,035369014 | 10,49655466 |
| rh_R_STSda | female | B-Norm | 19,7429093 | 6,102572361 |
| rh_R_p24pr | female | B-Norm | 0,929161114 | 10,36833111 |

### 4. Supplementary information to results section: Association with puberty scores

| 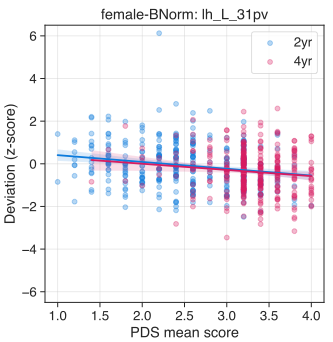 | 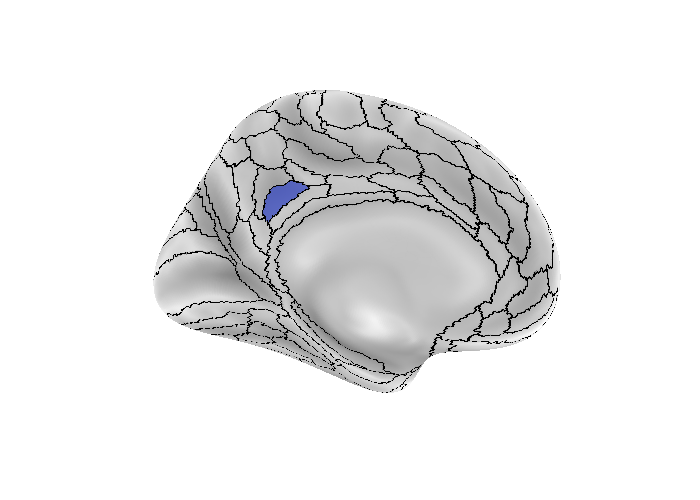 |
| --- | --- |
| Figure S 23 Association experiment: PDS with z - C-Norm.  Area 31pv of the left hemisphere yielded a significant association effect at the 2-year follow-up (t=-3.985, p<0.0244) between mean PDS scores and the z (i.e., deviation scores) as obtained from the female C-Norm model. Left panel shows the scatter plot with regression lines. Colors correspond to either the 2-year (blue) or 4-year (red) data. Right panel shows the location of area 31pv. | |

#### 4.1. Adjusting for family structure in training- and test-set.

We performed the same analysis as in the main text, however adjusting for family relationships before training the C- and B-Norm models. We opted for this auxillary analysis because the ABCD dataset has high proportions of siblings/twins.

Before training, we removed all participants from the training set that had a familial relationship with participants in the test set. If there were family relations in the test set, we randomly chose one of the siblings to remain in the test set. This resulted in 37 and 26 less males (new N=402) and females (new N=340) respectively.

Interestingly, we did not find the previously reported associations in the reported areas (see main text). Significance values increased to 0.0965 for the left 31pv and to 0.0636 for the left LO2, IFSa, PoI2 and LO3.

Given these results we briefly investigated the “family-adjusted” models for performance and difference in predicted norms compared to the “all-subs” models. We further tested whether the statistics (e.g., p-values) for the association analysis (“PDS ~ z”) was markedly different between the “all-subs” and “family-adjusted” models.

4.1.1. Differences in performance

Similar to the analysis of performance measures in the main text we investigated whether adjusting for family structure leads to differences in performance measures. Looking at the similar structures in the graphs below, we opted to not do the extra statistics because the patterns between the “all-subs” and “family-adjusted” models appears largely identical.

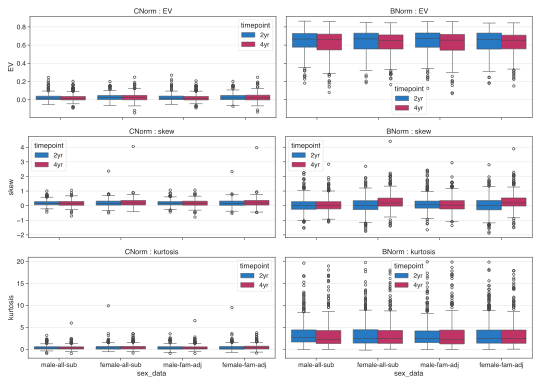

Figure S 24. Comparing performance metrics between C- and B-Norm models using all-subs or family-adjusted data.

Each panel depicts one of three performance measures for the models using all- or family-adjusted subjects (denoted as <sex>-all-sub or <sex>-fam-adj on the x-axis respectively). From top to bottom: explained variance (EV), skew, and kurtosis for both models (columns).As is evident from the pattern of the box plots, we do not see any marked effect of adjusting for family relationships in the training or test set.

4.1.2. Differences in predicted norms

We evaluated whether adjusting for family-relationship yields differences between the norms within each of the C- and B-Norm models. To do so, we loaded the predicted values (yhat) of the test set and used the PCNtoolkit to warp them (since our model uses the SinArcsin warp; see Methods in the main text).

For each model (C-Norm and B-Norm), we computed the normalized Euclidean distance between the predicted values obtained using the all-subject data and those obtained using the family-adjusted data.We repeated this process for all 360 Glasser regions.

Normalized Euclidean Distance values indicated high overlap between C- and B-Norm model norms suggesting that adjusting for family structure had likely no effect on model parameters (see table below).

Table S 10. Normalized Euclidean Distance between predicted norms of the test set using models trained with all participants and those trained adjusting for family structure.

|  |  | **Normalized Euclidean Distance** | | | |
| --- | --- | --- | --- | --- | --- |
| **Sex** | **Model** | **Min** | **Max** | **Mean (across ROIs)** | **Std**  **(across ROIs)** |
| **Male** | C-Norm | 0.00241 | 0.00984 | 0.00079 | 0.00069 |
|  | B-Norm | 0.00014 | 0.01117 | 0.00061 | 0.00075 |
| **female** | C-Norm | 0.00020 | 0.06120 | 0.00098 | 0.00331 |
|  | B-Norm | 0.00013 | 0.02385 | 0.00064 | 0.00129 |

4.1.3. Potential sample size problem?

We performed a correlation analysis to assess whether a drop in sample sizes potentially affected our significant ROIs. We used Pearson's correlation across all ROIs of -log10-transformed p-values for each of the conditions (i.e., for each sex specific model for each timepoint) of the "all-subs" test set and the "family-adjusted" test set. The correlations can be seen in the table below.

Interestingly, we observed lowes correlations for the 2-year follow-up data for male B-Norm models. Importantly, the correlations for the female B-Norm models of the 4-year follow-up data (which showed most marginally significant results) were almost perfectly correlated.

Table S 11. Pearson’s correlation of p-values from “PDS~z” GLM association tests between models using all participants (as determined by preprocessing) and models trained with family-adjusted data..

| **Sex** | **Model** | **Timepoint** | **Pearson’s rho** |
| --- | --- | --- | --- |
| *male* | *CNorm* | *2yr* | 0.877 |
|  |  | *4yr* | 0.914 |
|  | *BNorm* | *2yr* | 0.778 |
|  |  | *4yr* | 0.934 |
| *female* | *CNorm* | *2yr* | 0.960 |
|  |  | *4yr* | 0.959 |
|  | *Bnorm* | *2yr* | 0.955 |
|  |  | *4yr* | 0.979 |

### 5. Validation of normative models using percentile shifts

As a final validation step, similar to longitudinal changes used in pediatrics, we validated our normative models by categorizing participants into three groups based on percentile shifts between the 2-year and 4-year follow-up data: negative (zDiff < -1), stable (-1 < zDiff < 1), and positive (zDiff > 1), with zDiff representing the change in ROI-specific z-scores over time. For each ROI, participants were assigned to one of these groups, and a Kruskal-Wallis test was conducted to assess group differences in ΔPDS scores (4-year minus 2-year). No group differences survived multiple comparison correction using the Benjamini-Hochberg procedure (male: min p-uncorrected=0.001142, p-corrected=0.2723 for right PFop in the B-Norms; female: min p-uncorrected=0.001286, p-corrected=0.4526 for left PHA2 in the C-Norms). Nevertheless, regions showing uncorrected p-values < 0.05 are displayed in **Figure 4** for the female C-Norms (left) and B-Norms (right), along with corresponding boxplots of delta PDS scores for the most notable ROIs (TGv and PHA2 in the C-Norms; p24pr and PHA2 in the B-Norms, all left hemisphere). Of note, due to insufficient sample sizes within deviation groups, Kruskal-Wallis tests could not be performed in the following regions of the female C-Norms: MT, VMV1, and L31a (left hemisphere), and TE1a, PHT, PH, TPOJ1, VMV1, VMV2, and TE1m (right hemisphere). Figure S24 shows a similar graph for the male models.

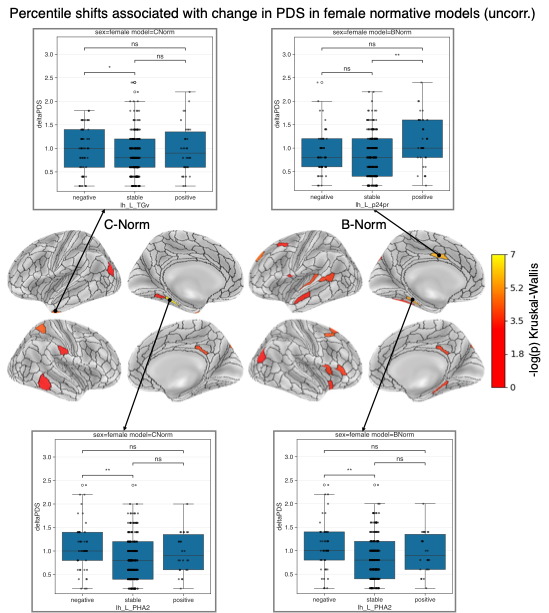

Figure S 25 Percentile shifts as indicators of individual level brain development and pubertal progress

Uncorrected significant -log-transformed p-values of a Kruskal-Wallis test plotted on the surface brain for female C-Norms (left) and B-Norms (right). Arrows connecting ROIs with boxplots indicate the two largest effects for an area in the left temporal pole (TGv) and left peri-hippocampal area PHA2 for the C-Norms or for parts of the left cingulate cortex (area p24pr) and area PHA2 for the B-Norms. The boxplots show delta PDS scores across the three groups: negative (zDiff < -1), stable (-1 < zDiff < 1), and positive (zDiff > 1). Black dots represent individual participants. Asterisks above connecting boxes indicate the significance of pair-wise Dunn’s tests (n.s.: p>0.05, *: p<0.05, **: p<0.01).

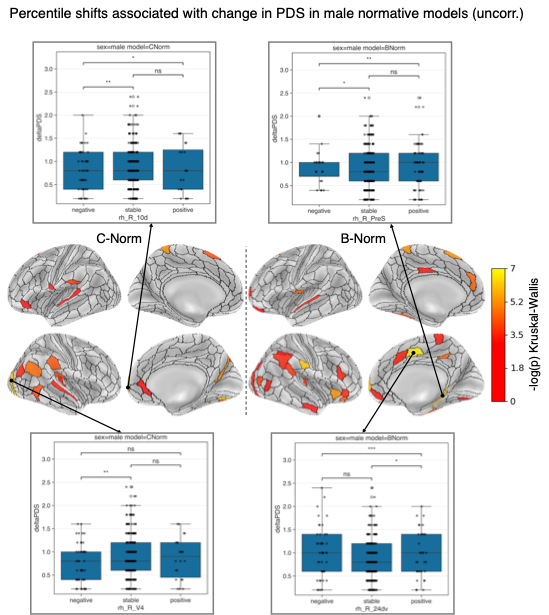

Figure S 26 Validation of male normative models with percentile shifts and change in pubertal progress.

Uncorrected significant -log-transformed p-values of a Kruskal-Wallis test plotted on the surface brain for male C-Norms (left) and B-Norms (right). Arrows connecting ROIs with boxplots indicate the two largest effects for an area in the right visual (V4) and frontal cortex (10d) for the C-Norms or for a section of the left frontal cortex (area p24pr) and the presubiculum (PreS) for the B-Norms. The boxplots show delta PDS scores across the three groups: negative (zDiff < -1), stable (-1 < zDiff < 1), and positive (zDiff > 1). Black dots represent individual participants. Asterisks above connecting boxes indicate the significance of pair-wise Dunn’s tests (n.s.: p>0.05, *: p<0.05, **: p<0.01).
